# Tuberculosis alters immune-metabolic pathways resulting in perturbed IL-1 responses

**DOI:** 10.1101/2020.12.17.423082

**Authors:** Alba Llibre, Nikaïa Smith, Vincent Rouilly, Munyaradzi Musvosvi, Elisa Nemes, Céline Posseme, Simba Mabwe, Bruno Charbit, Stanley Kimbung Mbandi, Elizabeth Filander, Hadn Africa, Violaine Saint-André, Vincent Bondet, Pierre Bost, Humphrey Mulenga, Nicole Bilek, Matthew L Albert, Thomas J Scriba, Darragh Duffy

## Abstract

Tuberculosis (TB) remains a major public health problem with host-directed therapeutics offering potential as novel treatment strategies. However, their successful development still requires a comprehensive understanding of how *Mycobacterium tuberculosis* (*M.tb*) infection impacts immune responses. To address this challenge, we applied standardised immunomonitoring tools to compare TB antigen, BCG and IL-1β induced immune responses between individuals with latent *M.tb* infection (LTBI) and active TB disease, at diagnosis and after cure. This revealed distinct responses between TB and LTBI groups at transcriptomic, proteomic and metabolomic levels. At baseline, we identified pregnane steroids and the PPARγ pathway as new immune-metabolic drivers of elevated plasma IL-1ra in TB. We also observed dysregulated induced IL-1 responses after BCG stimulation in TB patients. Elevated IL-1 antagonist responses were explained by upstream differences in TNF responses, while for IL-1 agonists it was due to downstream differences in granzyme mediated cleavage. Finally, the immune response to IL-1β driven signalling was also dramatically perturbed in TB disease but was completely restored after successful antibiotic treatment. This systems immunology approach improves our knowledge of how immune responses are altered during TB disease, and may support design of improved diagnostic, prophylactic and therapeutic tools.

## INTRODUCTION

As the leading cause of death by infection, tuberculosis (TB) is a major global public health problem^1^. It is estimated that one fourth of the world’s population is infected by its causative agent, *Mycobacterium tuberculosis* (*M.tb*). *M.tb* infection results in a diverse clinical spectrum, which includes asymptomatic latent infection (LTBI), incipient or subclinical stages, and active TB disease (TB), which occurs in 5-10% of infected persons^2,3^.

*M.tb* alters the host’s ability to clear infection by targeting immune and metabolic pathways. Initial research in mice, later supported by human studies^4^, demonstrated that interferon γ (IFNγ), tumour necrosis factor (TNF) and interleukin (IL)-1β are essential cytokines for immune control of *M.tb*^5–8^. Signalling through the IL-1 receptor can be regulated by binding of the IL-1 receptor antagonist (IL-1ra)^9^. Levels of circulating IL-1ra are elevated in TB patients, and have been proposed as a potential biomarker for diagnosis of active disease or response to treatment^10^. However, little is known about the underlying mechanisms that lead to higher IL-1ra concentrations in active TB disease. Metabolic reprogramming is also crucial for determining successful immune responses^11^. Furthermore, immune-metabolic signatures associated with TB progression have previously been described^12,13^.

To better understand how these crucial cytokine pathways are perturbed in active TB disease, we applied a standardised immunomonitoring tool^14,15^ to compare induced immune responses between LTBI and TB patients at proteomic, transcriptomic, and metabolomic levels. Relevant stimuli that were used included *M.tb* antigens (TB Ag), Bacillus Calmette-Guérin (BCG), IL-1β, and a Null control. This approach revealed multiple differences at the proteomic and transcriptomic levels between LTBI and TB patients. Integration of induced cytokine responses with baseline metabolic profiles helped to identify unique immune and metabolic drivers of elevated plasma IL-1ra in TB patients, in particular, the PPARγ pathway. After BCG stimulation, IL-1ra was secreted at higher levels in active TB patients compared to LTBI. Experimental analysis revealed that this was partly due to differences in TNF mediated signalling. Furthermore, IL-1α and IL-1β were secreted at lower levels in active TB patients due to differences in post-translational modifications. Finally, IL-1β stimulation revealed a dysregulated IL-1 signalling response in TB patients, which was restored after successful treatment. Overall, our approach adds new understanding of how *M.tb* impacts human immune responses, providing new avenues for better diagnostic tools, vaccines and treatments, including host-directed therapeutics.

## RESULTS

### Immune stimulation identifies specific gene expression differences that improve stratification of active TB disease from latent *M.tb* infection

To examine how TB disease perturbs immune responses, we stimulated whole blood from active TB patients and LTBI controls with relevant stimuli that included TB Ag, BCG and IL-1β, plus a non-stimulated control (Null). Principal component analysis (PCA) of 622 immune genes measured showed clustering of stimuli-specific responses, with 54% of the total variance captured by the first three principal components (Figure 1A and Suppl Figure 1A). As previously reported^16^, several genes were differentially expressed in active TB disease compared to LTBI in the absence of stimulation, namely 32% (200 genes, q <0.001) of the immune genes examined.

**Figure 1.**
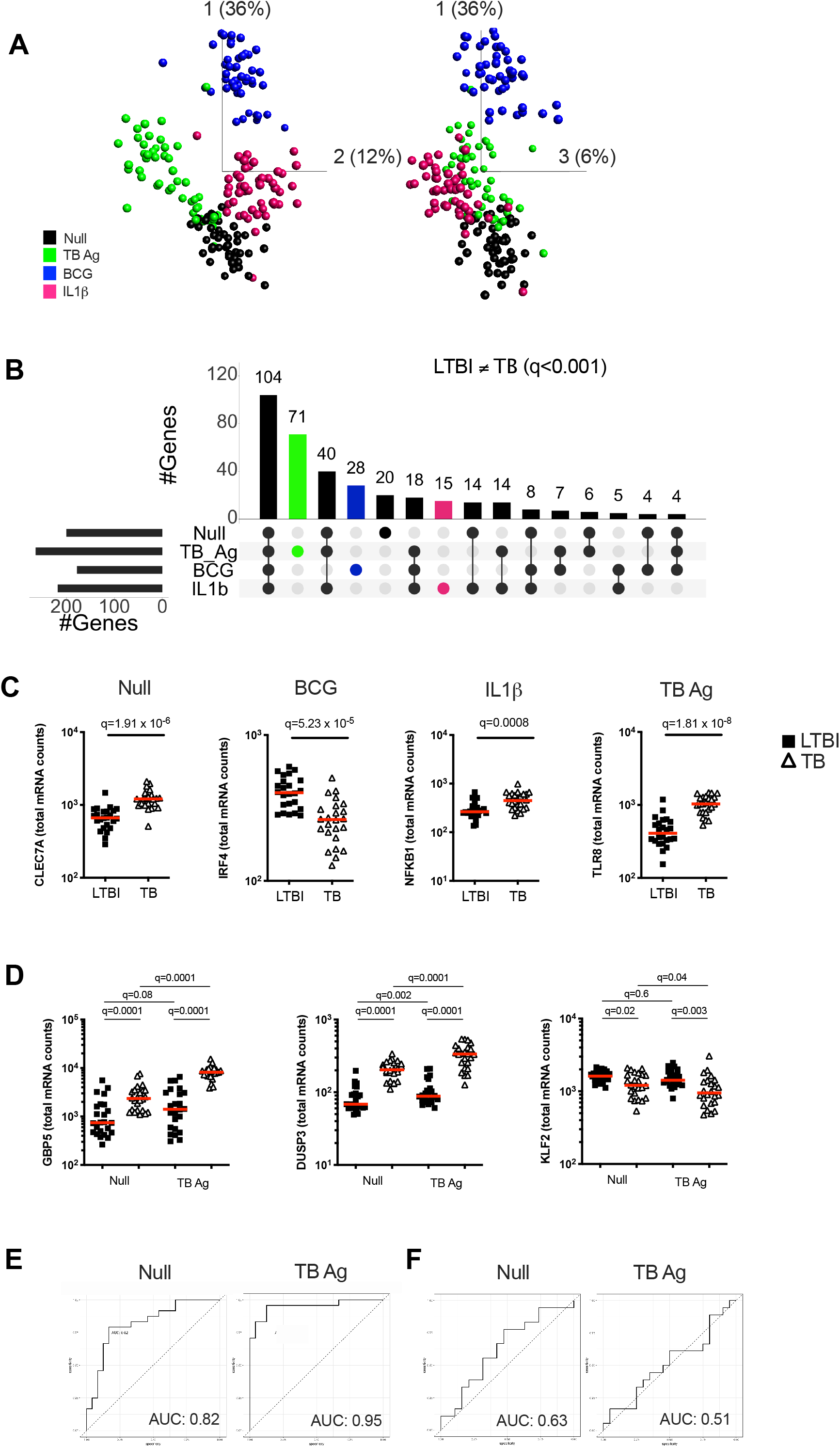
Immune stimulation identifies specific gene expression differences between latent *M.tb* infection and active TB disease. **(A)** Principal component analysis (PCA) on expression of 622 genes from 24 individuals with latent *M.tb* infection (LTBI) and 24 with active TB disease (TB) after whole blood stimulation with TB Ag, BCG, IL-1β, and a non-stimulated control (Null). Each coloured circle represents one individual for each condition. **(B)** Upset plot showing the intersection of differentially expressed genes (q value <0.001, Mann-Whitney corrected for multiple comparisons) between TB and LTBI, across the four stimulations. **(C)** Examples of significantly different stimuli-specific transcripts (q<0.001) between TB and LTBI. **(D)** Expression of the 3 genes from the Sweeney TB score (Sweeney3) in the Null and TB Ag conditions on visit 1 pre-treatment. Area under the receiver operating characteristic curve (AUC) for discrimination of LTBI/TB groups using Sweeney3 for the Null and TB Ag conditions on visit 1 pre-treatment (Pre-Tx) **(E)** and on visit 2 after successful antibiotic treatment (Post-Tx) **(F)**. Comparisons of LTBI/TB groups within the same stimulation were performed using unpaired non-parametric t-tests; comparisons between Null and stimulated conditions within the LTBI/TB groups were performed using a paired non-parametric t-test. Correction for multiple comparisons was then applied. Red line: median values. Closed square: LTBI, Open triangle: TB.

Focusing on differential genes that were revealed only by immune stimulation, we compared LTBI and TB groups using a t-test with a stringent cut-off value of q<0.001. This resulted in 114 genes (18% of total) with differences between LTBI and TB groups only after specific stimulation (TB Ag; 71, BCG; 28, IL-1β; 15) (Figure 1B and Supplementary Table 2), with TB Ag stimulation revealing significantly more differences than the other stimulations (Supplementary Table 3). Examples of stimuli-specific induced differences for each condition and the Null are illustrated in Figure 1C, Supplementary Figure 1B and Supplementary Table 4.

Whole blood transcriptomic approaches, based on signatures that primarily comprise interferon stimulated genes (ISG), are being developed for TB diagnosis, with promising strategies based on a 3-gene signature (Sweeney3 signature) in which a TB score is calculated using the formula (GBP5 + DUSP3)/2 – KLF2^17^, or the 11-gene blood TB risk signature, RISK11^18^. As TB Ag stimulation induced a high number of differences between LTBI and TB patients, we tested whether standardised immune stimulation could improve the performance of these signatures by comparing ROC curve analysis from Null and TB Ag stimulated samples (Figure 1D and 1E; Suppl Figure 1D and 1E). For the Sweeney3 signature, TB Ag stimulation accentuated the pre-existing differences at baseline, which translated into a superior area under the receiver operating characteristic curve (AUC) for Null_AUC_ of 0.82 [95% CI (0.66-0.94)] and for TB Ag_AUC_ of 0.95 [95% CI (0.88-1)] improving the ability to classify LTBI and TB cases (p=0.02) (Figure 1E). Similar results were obtained with the RISK11 signature, with Null_AUC_ of 0.82 [95% CI (0.69-0.94)] and TB Ag_AUC_ of 0.97 [95% CI (0.93-1)] (p=0.02) (Suppl Figure 1E). These differences were not present after successful antibiotic treatment of the TB patients in the Sweeney3 signature (Figure 1F; Null_AUC_ = 0.63 [95% CI (0.45-0.82)] and TB Ag_AUC_ = 0.51 [95% CI (0.32-0.71)], (p=0.4) or the RISK11 signature (Suppl Figure 1F; Null_AUC_ = 0.57 [95% CI (0.38-0.77)] and TB Ag_AUC_ = 0.63 [95% CI (0.45-0.81)] (p=0.7), confirming the specificity of these signatures.

In summary, immune stimulation revealed multiple differences between latent *M.tb* infection and active TB disease that likely reflect relative differences in ISG expression in blood leukocytes, which may help to better understand perturbed immunity during TB pathogenesis and potentially improve new diagnostic strategies.

### The IL-1 pathway is dysregulated in active TB disease

The TB Ag stimulation illustrated in Figure 1D and 1E captures differences in antigen-specific responses of CD4+ T cells, and subsequent downstream responses, between LTBI and TB patients. To explore the broader impact of TB disease in mounting efficient immune responses to complex stimuli, we investigated the response to BCG. To identify immune pathway differences, we performed correlation analysis between BCG-induced cytokines at the protein level (FC>1.3) in the LTBI and TB groups (Figure 2A). This revealed negative correlations in the control LTBI group, that were absent or markedly weaker in TB patients, suggesting altered regulation of cytokine production in TB disease. This included several key cytokines important for immune responses against TB such as IL-1α, IL-1β, IL-10, GM-CSF (CSF2) and TNF that all showed a positive correlation with the IL-1 receptor antagonist (IL-1ra) in TB patients, but not in LTBI (Figure 2A).

**Figure 2.**
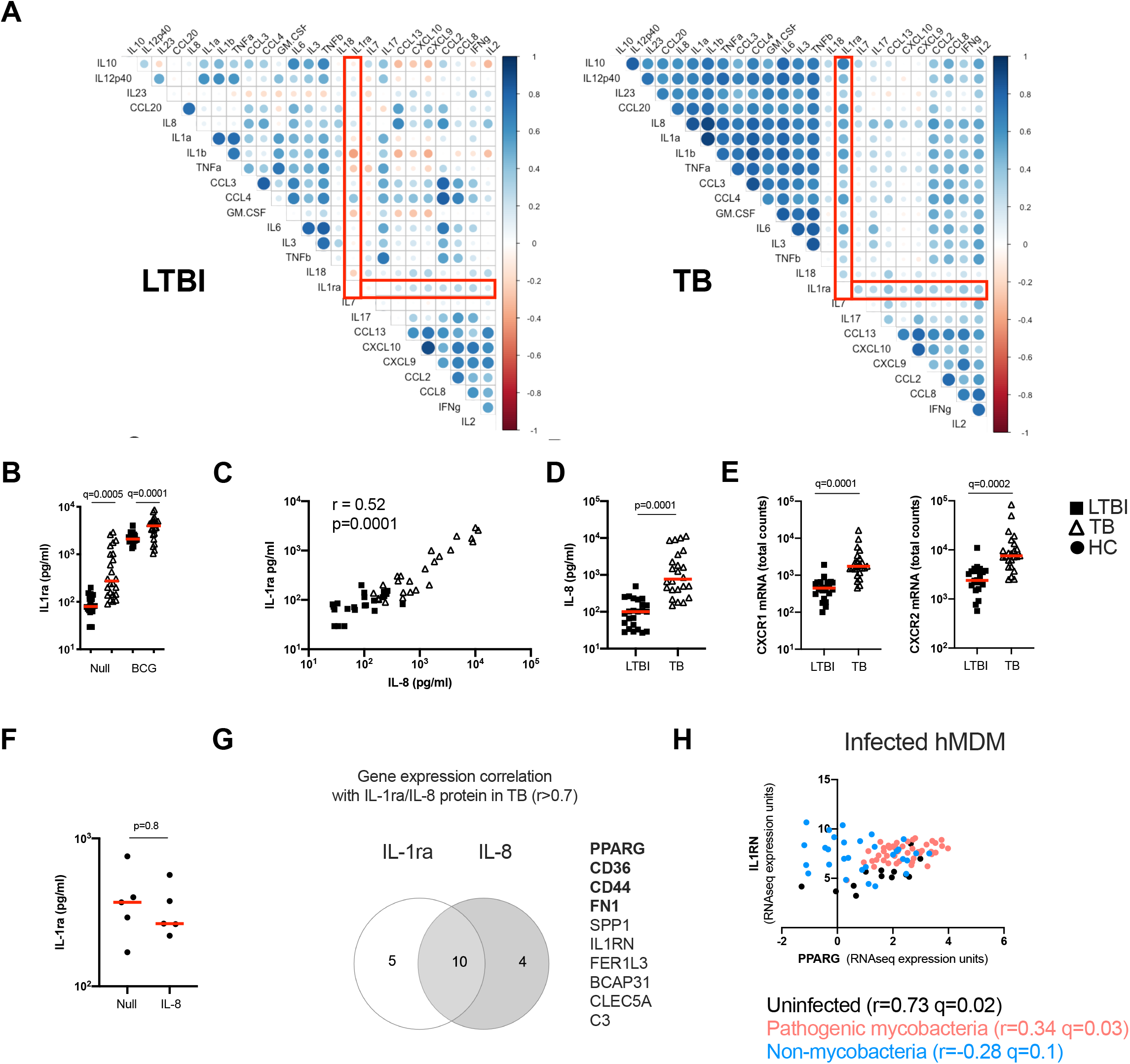
The IL-1 pathway is dysregulated in active TB disease. **(A)** Spearman correlation plot of the induced (Fold-change 1.3) proteins measured by Luminex in TruCulture supernatants after BCG stimulation for LTBI and TB. Red rectangles highlight IL-1ra correlations. **(B)** Levels of IL-1ra protein measured by Luminex in the Null and BCG conditions for LTBI and TB. **(C)** Spearman correlation between IL-8 and IL-1ra protein levels measured by Luminex in the Null condition. **(D)** IL-8 concentration measured by Luminex in the Null condition, for LTBI and TB. **(E)** Gene expression levels of the IL-8 receptor subunits CXCR1 and CXCR2 measured by Nanostring at baseline. **(F)** IL-1ra protein concentration measured by ELISA in supernatants of whole blood from healthy individuals stimulated with IL-8. **(G)** Venn diagram of the genes correlated with IL-1ra and IL-8 in the Null condition in TB, measured by Nanostring and Luminex, respectively. Genes associated with the PPARγ pathway are in bold. **(H)** Reanalysis of RNAseq data by Spearman correlation between PPARG and IL1RN gene expression levels of human monocyte-derived macrophages infected with pathogenic mycobacteria and non-mycobacteria species for 18 and 48h^20^. Comparisons between groups were performed using unpaired or paired (F) non-parametric t-tests and correction for multiple comparisons was applied. Red line: median values. Closed square: LTBI, Open triangle: TB, Closed circle: Healthy Control (HC).

Because IL-1 signalling is essential for *M.tb* control^8,19^, we further explored differences within the IL-1 family members between LTBI and TB. We started by investigating levels of IL-1ra and observed that TB patients presented higher concentrations of IL-1ra both at baseline and after BCG stimulation (Figure 2B). To better understand the drivers of IL-1ra secretion at baseline in TB, we performed correlation analysis of the protein data comprising 40 cytokines/chemokines in the Null tube and showed that the strongest association (r=0.52, p=0.0001) was with IL-8 (Figure 2C), a cytokine which also had significantly higher levels in TB patients (Figure 2D). Expression levels of both IL-8 receptors (CXCR1 and CXCR2) were also higher in TB patients (Figure 2E), however stimulation of blood from healthy controls with IL-8 did not induce IL-1ra (Figure 2F). We next explored gene expression signatures commonly associated with these two cytokines and found genes related to the peroxisome proliferator-activated receptor (PPAR)γ pathway (Figure 2G). We validated our findings using a published dataset of transcriptional analysis of human monocyte-derived macrophages (hMDMs) infected with different bacterial strains^20^. This showed a weak, yet significant positive correlation (r=0.34, q=0.03) between IL1RN (the transcript for IL-1ra) and PPARG transcripts in hMDMs infected with pathogenic mycobacteria, but not with non-mycobacterial species (r= −0.28, q=0.1) (Figure 2H). These results identify an immuno-metabolic link between the PPARγ pathway and the receptor antagonist IL-1ra.

### Pregnane steroids activate the PPARγ pathway resulting in increased IL-1ra secretion in active TB

PPARγ is a ligand-activated transcription factor that plays an essential role in metabolism and energy homeostasis, particularly in adipogenesis^21,22^. To explore potential metabolic pathways involved in IL-1ra secretion, we performed unsupervised mass spectrometry analysis on supernatants from unstimulated LTBI and TB patients’ blood and examined which metabolites correlated with IL-1ra protein at baseline in the TB group. Interestingly, 5 of the top 6 ranked metabolites showing significant correlations (r>0.6, q≤0.05) with IL-1ra in TB patients belonged to pregnane steroid derivatives (Figure 3A and Supplementary Table 5). These correlations were observed in TB patients, and not in LTBI (Figure 3A and 3B). Interestingly, levels of these steroids did not differ between the LTBI and TB groups (Suppl Fig 2A), in contrast to PPARG expression, which was higher in TB (Suppl Fig 2B). Pregnane steroids have been shown to be potential ligands of PPARγ^23^, and PPARγ is mainly expressed by CD14+ monocytes (Human Cell Atlas). To investigate whether activation of PPARγ could drive IL-1ra secretion, we incubated CD14+ monocytes isolated from healthy donors, with combinations of the PPARγ agonist rosiglitazone and the PPARγ antagonist GW9662, in the presence of BCG. We validated our approach by measuring surface expression of CD36, a fatty acid translocase that is up-regulated by PPARγ^24–26^. CD14+ monocytes stimulated with rosiglitazone upregulated CD36, and this could be prevented with GW9662 (Suppl Fig 2C). We observed that PPARγ activation through rosiglitazone increased IL-1ra secretion by CD14+ monocytes, which could be inhibited by GW9662 (Figure 3C). To further confirm the role of PPARγ in IL-1ra secretion, we silenced PPARγ using small interfering RNA (siRNA). We confirmed PPARγ knockdown in CD14+ monocytes by measuring levels of PPARγ protein by flow cytometry (Suppl Fig 2D). Both at baseline and after BCG stimulation, silencing of PPARγ resulted in decreased IL-1ra secretion (Figure 3D), with this knockdown impacting both conditions equally (Figure 3E). This suggests that the PPARγ pathway is not involved specifically in BCG-induced IL-1ra, although these data confirm PPARγ as a key regulator of baseline IL-1ra production in TB disease.

**Figure 3.**
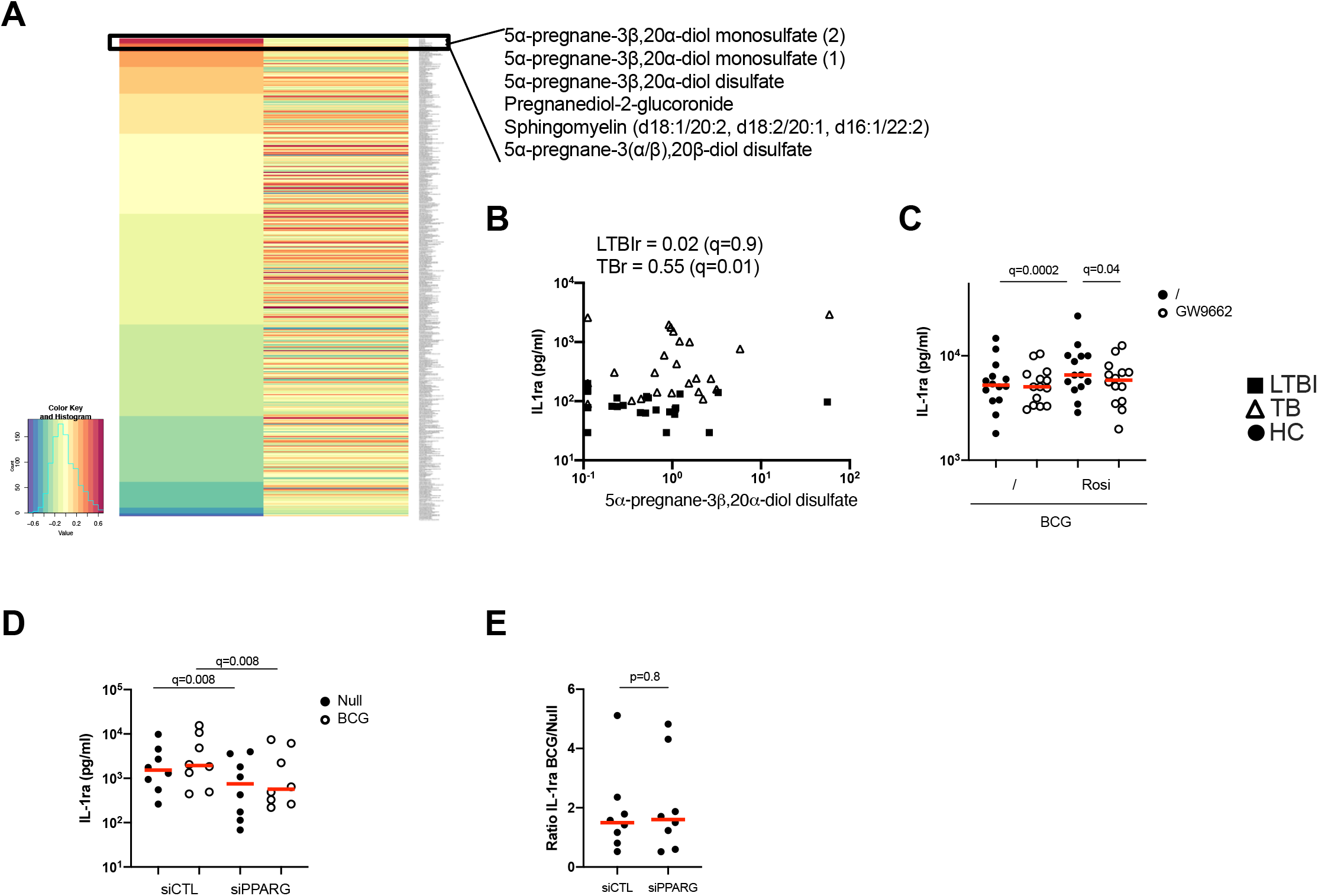
Pregnane steroids activate the PPARγ pathway resulting in increased IL-1ra secretion in active TB. **(A)** Heatmap of Pearson correlation coefficients between metabolites and IL-1ra protein in the Null condition, measured by mass spectrometry and Luminex, respectively, for TB (left) and LTBI (right). Metabolites are ordered in decreasing values of TB cases Pearson correlation coefficients. **(B)** Pearson correlation between IL-1ra levels measured by Luminex and 5α-pregnane-3β,20α-diol disulfate measured by mass spectrometry for LTBI and TB. **(C)** IL-1ra protein concentration measured by ELISA after stimulation of CD14+ monocytes from healthy individuals with BCG in the presence or absence of the PPARγ agonist rosiglitazone (Rosi) and/or the PPARγ antagonist GW9662. **(D)** IL-1ra protein concentration measured by ELISA after stimulation with BCG of PPARG silenced or control CD14+ monocytes from healthy individuals. **(E)** Ratio of IL-1ra induced upon BCG stimulation versus the Null Control in PPARG silenced or control CD14+ monocytes. Comparisons between groups were performed using paired non-parametric t-tests and correction for multiple comparisons was applied. Red line: median values. Closed square: LTBI, Open triangle: TB, Circle: Healthy Control (HC).

### Increased TNF signaling promotes IL-1ra secretion

Protein levels of IL-1ra were higher in TB patients both at baseline and after BCG stimulation (Figure 2B), the latter of which was not explained by PPARγ (Figure 3E). Therefore, we investigated the drivers of IL-1ra secretion in the context of BCG stimulation. Previous studies have identified TNF, IL-1 and type I and II IFNs as drivers of IL-1ra secretion^27–32^. To test this in our system we stimulated whole blood from healthy individuals with TNF, IL-1β, IFNγ, IFNβ, IFNα, and IL-8 and confirmed TNF and IFNβ (q=0.001) as the strongest inducers of IL-1ra (Figure 4A). Furthermore, TNF and type I IFNs acted synergistically in inducing IL-1ra, as the combination of these cytokines (TNF + IFNα/β) resulted in a 4-fold higher induction as compared to stimulation with each cytokine alone (Figure 4B). We then investigated the effect of blocking IFN and TNF signalling through the use of blocking antibodies. Blocking of TNFR (q=0.001), but not IFNAR (q=0.7), resulted in significantly less IL-1ra production (Figure 4C). To obtain further insight into the potential contribution of TNF and type I IFN in IL-1ra production upon BCG stimulation, we examined a previously published kinetic study of BCG stimulated whole blood^33^. This consisted of five healthy donors at 13 time points within 30h of BCG stimulation from which we measured IFNα and IFNβ to analyse with existing TNF and IL-1ra data. A time-dependent linear regression on IL-1ra using IFNα, IFNβ and TNF protein secretion as predictors revealed a strong association with TNF (q=0.002) but not type I IFNs (IFNα q=0.6, IFNβ q=0.6) (Figure 4D). IFNβ was not induced upon BCG stimulation, and IFNα was induced by BCG from the 10h time point (Figure 4D).

**Figure 4.**
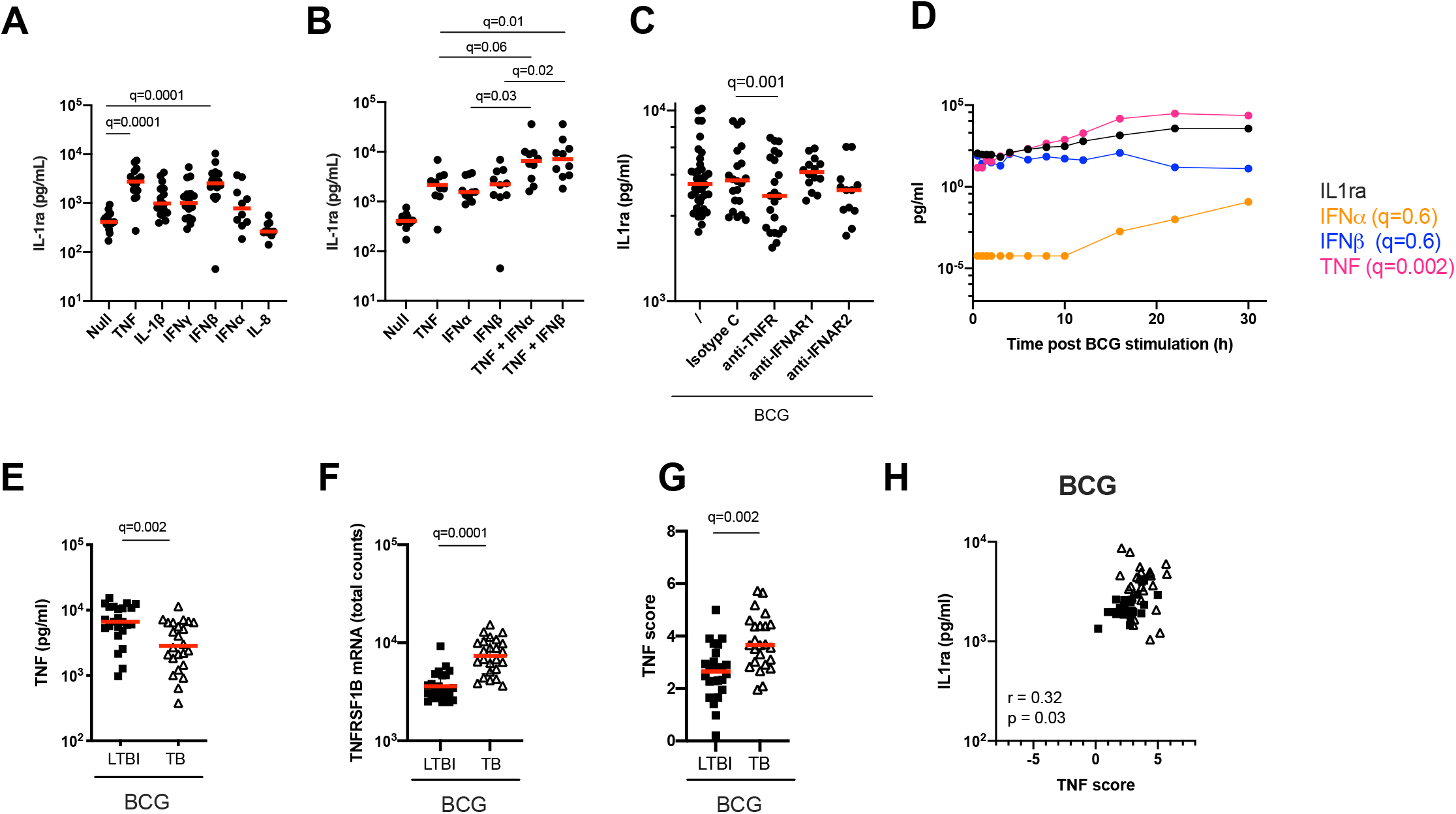
Increased TNF signaling promotes IL-1ra secretion. **(A)** Concentrations of IL-1ra measured by Luminex after stimulation of blood from healthy individuals with TNF, IL-1β, IFNγ, IFNα, IFNβ, and IL-8. **(B)** IL-1ra protein concentration measured by ELISA after whole blood stimulation with TNF, IFNα, IFNβ, TNF + IFNα and TNF + IFNβ. **(C)** IL-1ra protein concentration measured by ELISA after whole blood stimulation with BCG in the presence of anti-TNFR or anti-IFNAR, or their respective isotype controls. **(D)** Kinetics of IL-1ra, IFNα, IFNβ and TNF secretion measured by Luminex (IL-1ra and TNF) and Simoa (IFNα and IFNβ) upon BCG stimulation during the course of 30h. 1 representative donor of five healthy individuals is shown. **(E)** TNF protein concentration measured by Luminex in the BCG stimulated condition, for LTBI and TB. **(F)** TNFRSF1B mRNA expression levels measured by Nanostring in the BCG stimulated condition, for LTBI and TB. **(G)** TNF gene score in the BCG stimulated condition for the LTBI and TB groups. **(H)** Spearman correlation between IL-1ra levels and TNF gene score in the BCG stimulated condition. Paired (A, B, C) and unpaired (E, F and G) non-parametric t-test corrected for multiple comparisons. Red line: median values. Closed circle: Healthy Controls, Closed square: LTBI, Open triangle: TB.

To apply these findings to TB patients, we measured TNF protein after BCG stimulation and observed that levels of TNF were surprisingly higher in LTBI compared to TB (Figure 4E). We next examined expression levels of the TNF receptor subunit (TNFRSF1B) and observed significantly higher levels (q=0.0001) in TB patients (Figure 4F), which may reflect increased signalling. To test this hypothesis, we calculated a gene score based on unique cytokine-induced gene signatures (see Materials and Methods and Supplementary Table 6) as previously described^15^. The TNF gene score, which is a surrogate for quantifying signalling by this cytokine, was higher in the TB group compared to LTBI after BCG stimulation (Figure 4G), and also positively correlated (Rs=0.32, p=0.03) with IL-1ra protein concentrations (Figure 4H). These results demonstrate an essential role for TNF in BCG-induced IL-1ra, a response that is augmented in active TB disease.

### TB patients present lower granzyme concentrations which impact levels of functional IL-1α and IL-1β

BCG stimulation revealed dysregulation of multiple arms of the IL-1 pathway in active TB disease (Figure 2A). As well as observing differentially induced IL-1ra between LTBI and TB (Figure 2B), we also saw significant differences in IL-1α and IL-1β secretion (Figure 5A). In contrast to IL-1ra, both IL-1α and IL-1β protein were secreted at significantly higher levels in LTBI upon BCG stimulation. However, these differences were not mirrored at the transcript level (Figure 5B). We therefore hypothesized that post-transcriptional/post-translational modifications might be impaired during active TB disease. IL-1β requires post-translational cleavage to signal and engage functional outcomes, whereas IL-1α can manifest basal levels of activity in its unprocessed form^34^. As this protein processing can occur through either inflammasome-dependent or independent manners^34,35^, we explored whether both pathways were engaged in our whole blood stimulation system. We stimulated blood from healthy donors with BCG in combination with either an inflammasome inhibitor (the Syk inhibitor R406) or a serine-protease inhibitor (3,4-Dichloroisocoumarin, DCI) capable of blocking a range of enzymes involved in inflammasome-independent cleavage of IL-1 immature proteins, including granzymes and cathepsins. Both inhibitors resulted in a partial but significant reduction of IL-1β secretion after BCG stimulation (Figure 5C), supporting a role for the inflammasome and serine-proteases in the secretion of IL-1β in stimulated whole blood cultures.

**Figure 5.**
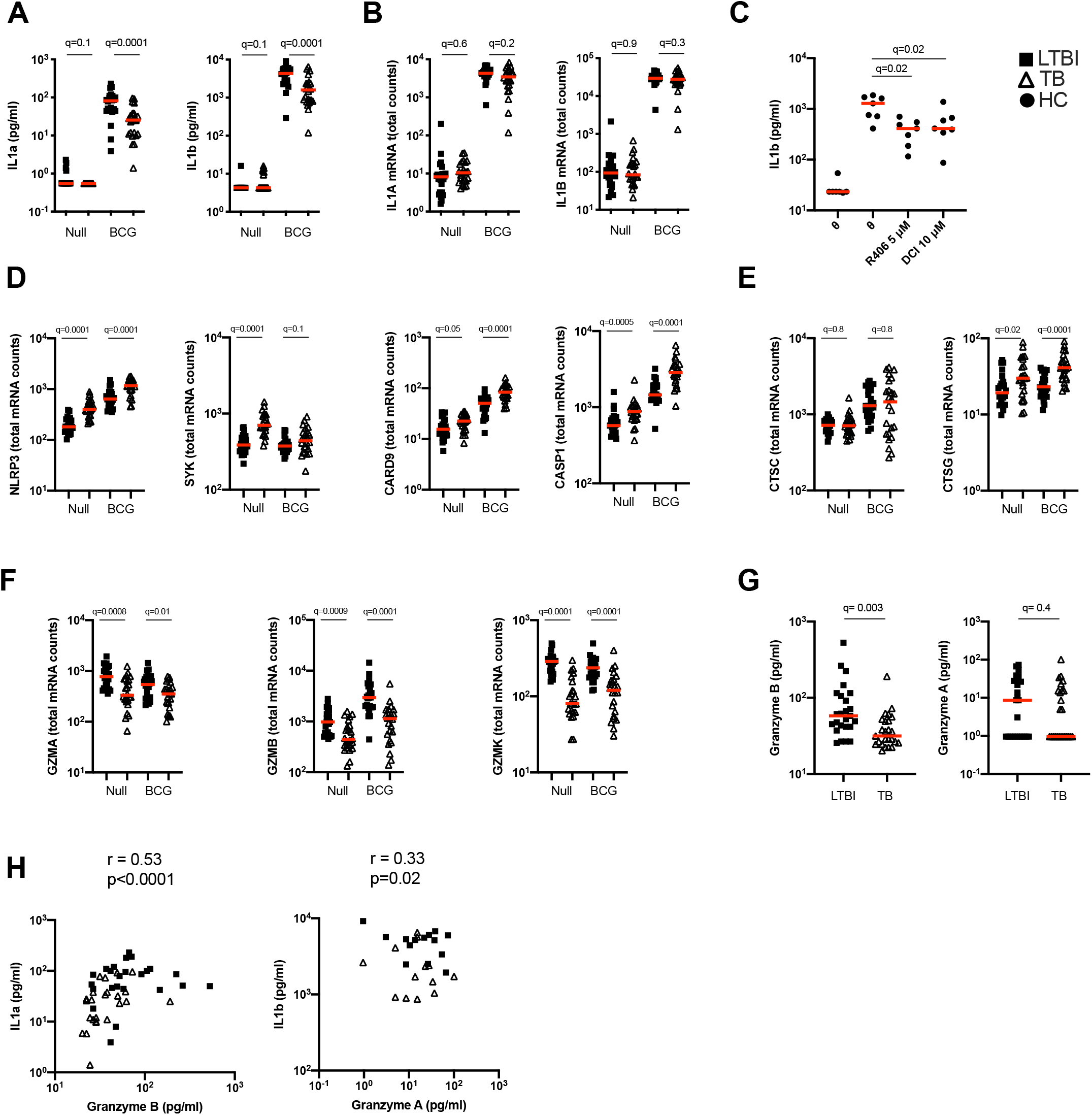
TB patients present lower granzyme concentrations which impact levels of functional IL-1α and IL-1β. **(A)** Levels of IL-1α and IL-1β protein measured by Luminex in the Null and BCG stimulated conditions. **(B)** Levels of IL1A and IL1B mRNA measured by Nanostring in the Null and BCG conditions. **(C)** BCG stimulation of healthy donor blood with the Syk inhibitor R406 (5 μM) or the serine-protease inhibitor 3,4-Dichloroisocoumarin (10 μM). Levels of mRNA expression for inflammasome components NLRP3, SYK, CRAD9 and CASP1 **(D)**, cathepsins **(E)** and granzymes **(F)** measured by Nanostring in the Null and BCG conditions for TB and LTBI. **(G)** Granzyme A and granzyme B protein concentrations in TruCulture BCG supernatants for TB and LTBI measured by Luminex. **(H)** Spearman correlation between protein concentrations of granzyme B and IL-1α, and granzyme A and IL-1β in BCG supernatants. Paired (C) and unpaired non-parametric t-test corrected for multiple comparisons. Red line: median values. Closed square: LTBI, Open triangle: TB, Closed circle: Healthy Controls.

We then examined differences in these pathways between LTBI and TB patients. The transcripts of inflammasome components NLRP3, SYK, CARD9 and CASP1 were significantly more expressed after BCG stimulation in TB patients compared to LTBI (Figure 5D). Cathepsin G, a neutrophil serine-protease that cleaves pro-IL-1β, was also expressed at higher levels in the TB group (Figure 5E), whereas no differences were observed in cathepsin C transcripts. In contrast, other genes that participate in inflammasome-independent IL-1 cleavage, such as granzymes A, B and K, were expressed at higher levels in the LTBI group (Figure 5F) suggesting that this pathway was more relevant for BCG induced IL-1β. Concentrations of granzyme B in the BCG supernatants were higher in the LTBI group compared to TB, however, no significant differences were observed with granzyme A (Figure 5G). It was notable that most granzyme A levels were below the limit of detection of the assay. Levels of granzyme B strongly correlated with IL-1α (R=0.53, p=0.0001) and there was a moderate but significant (R=0.33, p=0.02) correlation between IL-1β protein and Granzyme A (Figure 5H). These data support our hypothesis of aberrant IL-1 induction in TB disease, with lower levels of granzyme protein resulting in reduced cleavage of IL-1 precursors, and thus less active forms of IL-1 cytokines.

### IL-1β signalling is perturbed in active TB disease and is reset after successful antibiotic treatment

While BCG stimulation revealed perturbed secretion of IL-1 cytokines in active TB disease (Figure 2B and 5A), it did not allow examination of potential differences in IL-1 signalling. To investigate this, we stimulated whole blood from TB patients and LTBI controls with IL-1β and defined a signature of 107 induced genes (Null vs IL-1β q<0.001, Supplementary Table 7). Of this IL-1β driven gene signature, 72 genes were differentially expressed (q<0.001) between TB and LTBI (Figure 6A), with 95% of the differential genes showing higher levels of expression in TB than in LTBI. Among these differentially expressed genes were key signal transducers and transcription factors such as NKFB1/2 and IRAK2/3 (Figure 6B), suggesting altered downstream signalling. Analysis of 18 TB patients after successful antibiotic treatment showed that the perturbed IL-1β signalling response was completely reset and showed no differences compared to controls (Figure 6B, and Supplementary Table 8). Our data shows that IL-1β signalling is strongly perturbed in active TB disease, and this response is restored after successful treatment.

**Figure 6.**
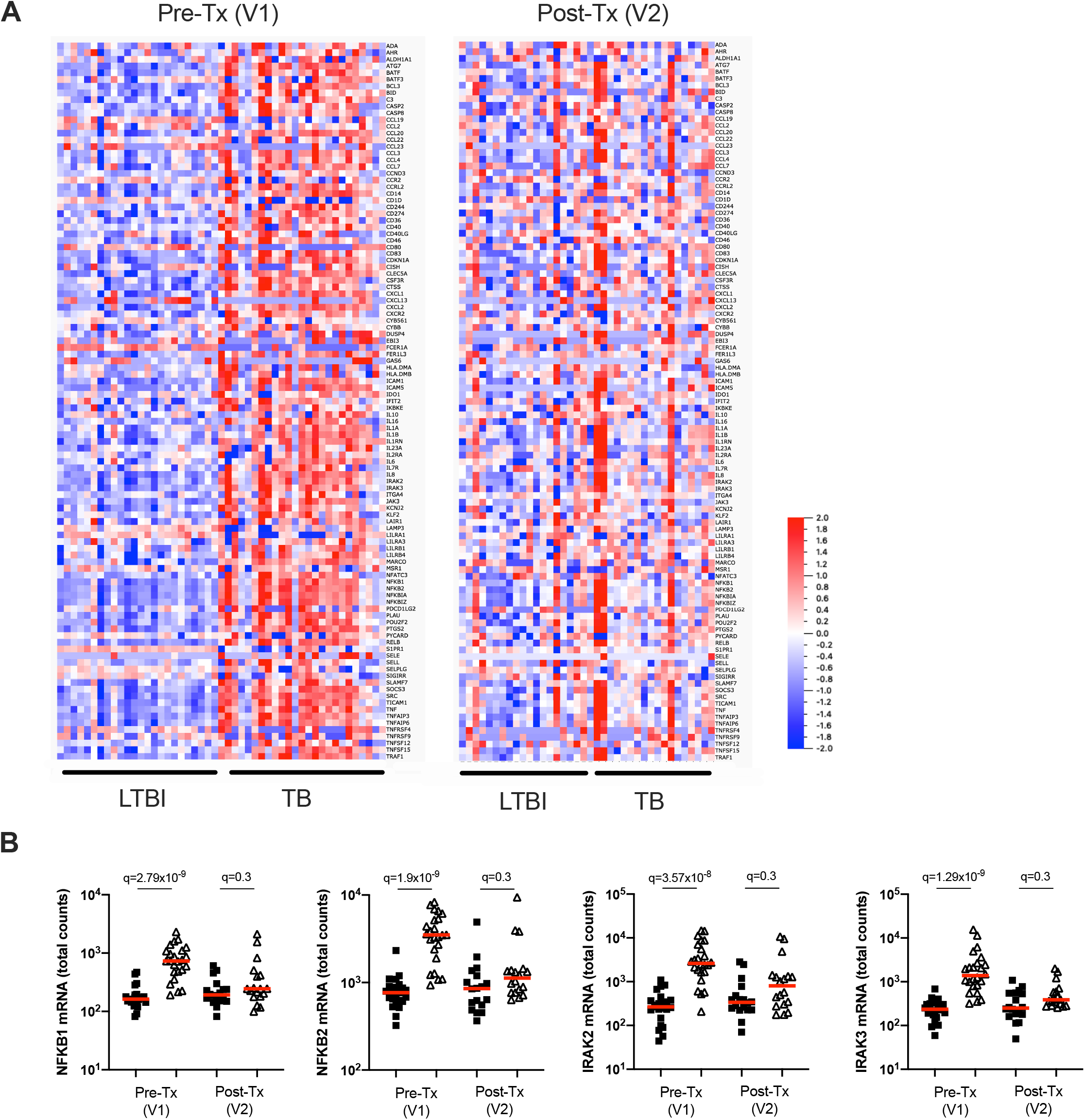
IL-1β signalling is perturbed in active TB disease and is reset after successful antibiotic treatment. **(A)** Heatmap showing expression levels of IL-1β-induced genes (Null vs IL-1β q=0.001) measured by Nanostring in LTBI and TB, in visit 1 pre-treatment (Pre-Tx V1) and visit 2 post-treatment (Post-Tx V2). **(B)** Levels of NFKB1, NFKB2, IRAK2 and IRAK3 mRNA measured by Nanostring after IL-1β stimulation for LTBI and TB. Unpaired non-parametric t-test corrected for multiple comparisons. Red line: median values. Closed square: LTBI, Open triangle: TB.

## DISCUSSION

A better understanding of how TB disease impacts immune responses is required for improved design of novel host-directed therapies. Here, we investigated differences in TB Ag, BCG and IL-1β induced immune responses between LTBI and TB patients using a standardised immunomonitoring system. Integration of transcriptomic, proteomic and metabolomic data sets at baseline and after immune stimulation revealed altered immune and metabolic pathways in TB disease. Previous studies have identified immune and metabolic differences in unstimulated whole blood of TB patients^36–38^. However, this approach is limited when studying impaired responses to immune challenges. Using TruCulture we identified and dissected how members of the IL-1 family, which are key cytokines for *M.tb* control, are perturbed in TB disease. The potential clinical relevance of such an approach was illustrated by the observation that immune stimulation improved the diagnostic score of a previously described gene expression signature for TB diagnosis^17^.

The antagonist IL-1ra molecule was elevated in TB at baseline conditions. Despite this cytokine having been proposed as a potential biomarker to distinguish LTBI from active TB^10^, the biological triggers of IL-1ra in the context of TB remain largely unknown. We showed that increased basal levels of IL-1ra in TB patients are associated with the PPARγ pathway. PPARγ is constitutively and highly expressed in healthy alveolar macrophages^39,40^, and has been shown to regulate host immune and metabolic responses to mycobacterial infection^41^. *M.tb*-driven activation of PPARγ promotes IL-8 secretion^42^, and activation of PPARγ in THP-1 cells results in IL-1ra secretion^43^, in agreement with our observations. Furthermore, Pott and colleagues identified IL1RN as a putative PPARγ target gene in a THP-1 model of human macrophages^44^. We propose a model in which pregnane steroids in TB disease activate the PPARγ pathway, which may act at different levels (eg. CD36, IL-1ra) to promote *M.tb* replication. *M.tb* can promote CD36 expression through PPARγ activation^45^, and CD36 contributes to *M.tb* survival^46,47^. Finally, the PPARγ axis can trigger IL-1ra secretion^43^, which inhibits IL-1 signalling, essential for pathogen clearance^8,19^. Inhibition of PPARγ has been associated with decreased mycobacterial burden in mouse and human macrophages^41,42,45^, and deletion of PPARγ in lung macrophages has been shown to be immunoprotective in the context of *M.tb* infection in mice^48^. PPARγ is therefore a good host-directed therapeutic target candidate, as it impacts lipid body biogenesis, cytokine production and *M.tb* replication^49^. PPARγ agonists are already being used in the treatment of conditions such as diabetes, where they act as insulin sensitizers^50^. Diabetes prevalence is escalating globally and the increasing overlap with TB is a major source of concern, with type 2 diabetic patients having a 3-fold increased risk of developing TB^51^. Activating PPARγ agents would be predicted to enhance susceptibility to TB disease. Therefore, careful assessment of how metabolic modulators impact comorbidities is required.

Stimulation with BCG revealed perturbed IL-1 family cytokine responses in TB patients, which is crucial for host control of *M.tb* infection^7,52,53^. Specifically, TB patients simultaneously had lower agonist (IL-1α and IL-1β) and higher antagonist (IL-1ra) responses. Detailed kinetic studies^33^, combined with single cytokine stimulations^15^, blocking experiments, and time series analysis permitted the identification of TNF as key driver of IL-1ra in TB disease. These results add to the existing complexities of cytokine responses to *M.tb,* in which the overall context will determine infection outcome, rather than the presence/absence of particular cytokines. For instance, the timing of IL-1^54,55^ or type I IFN signalling^56^ is a decisive factor to exert beneficial roles in the context of TB disease. Our study shows how TNF, a critical cytokine for control of *M.tb*, also induced IL-1ra expression, which can prevent effective IL-1 responses, that are in turn essential for protective immunity.

The observed IL-1α and IL-1β protein differences were not mirrored at the transcriptional level, suggesting defects in either post-transcriptional or post-translational regulation. Notably, both IL-1α and IL-1β require enzymatic post-translational cleavage, which can occur through inflammasome-dependent or – independent mechanisms^57^. *M.tb* has previously been shown to inhibit inflammasome activation and IL-1β processing^58^, and a recent study demonstrated that CARD9, an inflammasome protein associated with fungal induction of proinflammatory cytokines, negatively regulated IL-1β production during bacterial infection^59^. It has also been shown that IL-1β production in *M.tb* infection can take place independently of caspase-1 and ASC-containing inflammasomes in mice^53^. In our study we observed both higher inflammasome activity and lower granzyme expression in TB patients. Thus, lower granzyme activity in TB patients may result in lower post-translational cleavage of IL-1α/β. This is supported by previous studies in which PBMC stimulation with *M.tb*-related antigens resulted in lower levels of granzyme B in TB patients^60^. In addition, a recent multi-cohort study identified granzyme B, produced by polyfunctional NK cells, as a characterising feature of LTBI compared to uninfected controls^61^.

We and others have reported distinct secretion of IL-1 agonists and antagonists upon *M.tb* infection^8,10,62^, however, whether IL-1 signalling itself is impaired in TB disease remains poorly explored. Here, we showed perturbed IL-1β signalling in TB patients, with higher expression of most IL-1β-induced genes in TB compared to LTBI. This observation shows that there is no intrinsic defect in IL-1β signalling capabilities in TB patients. However the timing of IL-1 responses has been shown to be critical for *M.tb* control^54,55^, and elevated IL-1 signalling has been associated with lung immunopathology^63,64^. The consequences of IL-1β hyper responsiveness in TB patients therefore warrants further investigation. Importantly, the elevated IL-1β signalling was no longer present after successful antibiotic treatment, suggesting that immune responses were reset to healthy levels. A limitation of the current study is that experiments were performed on peripheral blood, whereas the cytokine networks that regulate *M.tb* infection *in vivo* also act on tissue resident cells, which might behave differently. Also, cellular differences between the groups studied, such as higher monocyte-to-lymphocyte and neutrophil-to-lymphocyte ratios in active TB compared to LTBI^65^, could potentially contribute to some of the observed differences.

In summary, our study of immune-metabolic responses revealed perturbed responses in TB disease at transcriptomic, proteomic and metabolomic levels. Integration of these datasets has improved our knowledge on how TB disease impacts immune-metabolic pathways resulting in perturbed IL-1 responses, which are essential for *M.tb* control. This better understanding of how *M.tb* alters immune responses may facilitate the improved design of preventive and therapeutic tools, including host-directed strategies.

## ACKNOWLEDGEMENTS

This study was funded by the Bill and Melinda Gates Foundation (OPP1114368 and OPP1204624), with additional support from the French Government’s Investissement d’Avenir Program, Laboratoire d’Excellence “Milieu Intérieur” Grant ANR-10-LABX-69-01. AL was supported by the Fondation Recherche Médicale (SPF20170938617) and the European Commision (H2020-MSCA-IF-2018, 841729). NS was supported by an Institut Pasteur Roux Cantarini fellowship. We thank the UTechS CB of the Center for Translational Research, Institut Pasteur for supporting Nanostring analysis. DD thanks Immunoqure for provision of the mAbs under an MTA for the Simoa IFN-α assay. We are grateful to the study participants and the SATVI clinical and laboratory teams.

## AUTHOR CONTRIBUTIONS

AL designed and performed experiments, analysed and interpreted data and wrote the manuscript. NS designed and performed experiments, and analysed and interpreted data. VR analysed data. MM and EN performed experiments on the TB patient cohort. MM, CP, SM, BC, SK, VSA, VB, PB, HM and NB performed specific experiments and/or analysis. EF and HA were the clinical and laboratory study coordinators in South Africa, respectively. MLA, TJS and DD conceived the whole study, obtained funding and provided overall guidance. DD designed and supervised the whole study, designed experiments, analysed and interpreted data and wrote the manuscript. All authors contributed to manuscript revision, read and approved the submitted version.

## DECLARATION OF INTERESTS

MLA is a current employee of Insitro, who had no influence on the study design or reporting. The other authors declare no competing interests.

## FIGURE LEGENDS

**Supplementary Figure 1. (A)** Scree plot showing the variance explained by each PCA component on the gene expression (Nanostring) dataset. **(B)** Examples of significantly different (q<0.001) stimuli-specific induced transcripts between TB and LTBI. **(C)** Expression of the 3 genes from Sweeney3^17^ in the Null and TB Ag conditions on visit 2 post-treatment. **(D)** RISK11 score^66^ in the Null and TB Ag conditions, pre and post-treatment, for the LTBI and TB groups, calculated using the Singscore method. Area under the receiver operating characteristic curve (AUC) for discrimination of LTBI/TB groups using the RISK11 score for the Null and TB Ag conditions on visit 1 pre-treatment (Pre-Tx) **(E)** and visit 2 after successful antibiotic treatment (Post-Tx) **(F)**. Comparisons of LTBI/TB groups within the same stimulation were performed using unpaired non-parametric t-tests; comparisons between Null and stimulated conditions within the LTBI/TB groups were performed using a paired non-parametric t-test. Correction for multiple comparisons was then applied. Red line: median values. Closed square: LTBI, Open triangle: TB.

**Supplementary Figure 2. (A)** Relative levels of pregnane steroids identified in Figure 3A for both LTBI and TB, measured by mass spectrometry in the Null condition. **(B)** Gene expression levels of PPARG in the Null condition measured by Nanostring both in LTBI and TB. **(C)** Protein levels of CD36 measured by flow cytometry on the surface of CD14+ monocytes from healthy individuals in the presence or absence of the PPARγ agonist rosiglitazone (Rosi) and/or the PPARγ antagonist GW9662. **(D)** Intracellular protein levels of PPARγ measured by flow cytometry in CD14+ monocytes from healthy individuals after treatment with PPARG small interfering RNA (siRNA) or control. Comparisons between groups were performed using unpaired (A, B) or paired (C) non-parametric t-tests and correction for multiple comparisons was applied. Red line: median values. Closed square: LTBI, Open triangle: TB, Circle: Healthy Control (HC).

## MATERIALS AND METHODS

### Participant groups

Adult patients with sputum Xpert MTB/RIF-positive TB disease who tested HIV-negative were identified and recruited at clinics in Worcester, South Africa^62^. Healthy QuantiFERON-TB Gold (QFT) positive (*M.tb-*infected) adults were recruited from communities living in or around Worcester and those who matched TB cases by age and gender were enrolled as LTBI controls (Table 1). Blood was collected prior to treatment initiation in active TB cases and again 12-18 months later, after successful completion of treatment defined by clinical cure (V2, n=18). For the QFT+ controls, blood was also collected at a second time-point, 12-18 months after the initial visit (V2, n=19). The TB clinical study protocols and informed consent forms were approved by the Human Research Ethics Committee of the University of Cape Town (ref: 234/2015). Healthy donor blood was obtained from the CoSImmGEn cohort of the Investigation Clinique et Accès aux Ressources Biologiques (ICAReB) platform, (Centre de Recherche Translationnelle, Institut Pasteur, Paris, France) or from the *Etablissement francais du sang* (EFS, France). Blood was collected in sodium-heparin tubes. Written informed consent was obtained from all study participants.

### TruCulture Whole Blood Stimulation

TruCulture tubes were prepared in batch as previously described^14,15,62^ with the following stimuli: QFT antigens (ESAT-6, CFP-10, TB7.7), Bacillus Calmette– Guérin (BCG; Sanofi Pasteur, 10^5^ units/ml), and IL-1β (Peprotec, 25 ng/ml). They were resuspended in 2 ml of buffered media and maintained at −20°C until use. Blood was collected in sodium-heparin tubes (50 IU/ml final concentration). Within 30 min of collection, 1 ml of whole blood was distributed into pre-warmed TruCulture tubes, inserted into a dry block incubator, and maintained at 37°C (+/− 1C), room air for 22 hr (+/− 15 min). After incubation, a valve was inserted to separate cells from the supernatant and to stop the stimulation reaction. Upon removal of the liquid supernatant, cell pellets were resuspended in 2 ml Trizol LS (Sigma), vortexed for 2 min, and rested for 10 min at room temperature before −80°C storage. Cytokine kinetic secretion data sets were previously described^33^. All stimulations were performed for 22h unless otherwise specified.

### Isolation and culture of blood leukocytes from healthy donors

For monocyte experiments, blood was collected in buffy coats from healthy donors sampled at the EFS, France. First, PBMCs were isolated by density centrifugation from peripheral blood leukocyte separation medium (Ficoll-Paque™ plus; GE Healthcare). Human monocytes were purified by negative selection with the classical monocyte isolation kit (Miltenyi). Monocytes were cultured at 1×10^6^ cells/mL in RPMI 1640 (Invitrogen, Gaithersburg, MD) (R10) containing 10% heat-inactivated fetal bovine serum, penicillin (100 U/ml), streptomycin (100 μg/ml; 1% pen-strep), and 1 mM glutamine (HyClone, Logan, UT).

### Protein analysis

Supernatants from TruCulture tubes were thawed on ice and tested by Luminex xMAP technology for a total of 32 proteins including cytokines, chemokines and growth factors as previously described^14^. Samples were measured on the Myriad RBM Inc platform (Austen, Texas, US) according to CLIA guidelines (set forth by the USA Clinical and Laboratory Standards Institute). The least detectable dose (LDD) for each assay was derived by averaging the values obtained from 200 runs with the matrix diluent and adding 3 standard deviations to the mean. The lower limit of quantification (LLOQ) was determined based on the standard curve for each assay and is the lowest concentration of an analyte in a sample that can be reliably detected and at which the total error meets CLIA requirements for laboratory accuracy. IL-1β, IL-1ra, IL-1α, IL-6, IP-10 and Granzyme A/B were also measured using Human ELISA Kit (ThermoFischer Scientific) or Human Custom Procarta-plexes (ThermoFischer Scientific), following manufacturer’s instructions.

To detect low concentrations of cytokines homebrew Simoa digital ELISA were used for IFNα^67^ and IFNβ^68^, as previously described.

### Gene expression analysis

Cell pellets in Trizol LS were thawed on ice for 60 min prior to processing. Tubes were vortexed twice for 5 min at 2000 rpm and centrifuged (3000 x g for 5 min at 4°C) to pellet the cellular debris generated during lysis. RNA was isolated from whole blood samples using the NucleoSpin 96 RNA tissue kit protocol (Macherey-Nagel) with some modifications as previously described^15^.

RNA integrity was assessed (Agilent RNA kits for the 2100 Bioanalyzer System). The NanoString nCounter system was used for the digital counting of transcripts. RNA was quantified using the Qubit RNA HS Assay Kit (Thermo Fischer Scientific) and 100ng of total RNA hybridized with the Human Immunology v2 (plus 30 additional genes (Supplementary Table 1) relevant for TB that were included) Gene Expression CodeSet according to the manufacturer’s instructions. Samples were processed in 7 batches (4 for Visit 1, 3 for Visit 2), within which the samples were randomized, and the same lot of reagents was used for all samples. All samples were normalized together following background subtraction of the negative control probes, using positive control probes and housekeeping genes (SDHA, HPRT1, POLR2A, RPL19, G6PD, TBP) selected by the GNorm method as previously described^69^. This was done using the nSolver™ analysis software (NanoString technologies). Quality control for our data involved checking the following metrics: fields of view counted (flag if < 0.75), binding density (flag if not in the 0.05-2.75 range), linearity of positive controls (flag if R2 < 0.9) and limit of detection for positive controls (flag if 0.5 fM positive control < 2 SDs above the mean of the negative controls).

### Metabolomic analysis

Supernatants from TruCulture Null tubes were thawed and tested by Ultrahigh Performance Liquid Chromatography-Tandem Mass Spectroscopy (UPLC-MS/MS) for a total of 696 metabolites. Samples were measured on the Metabolon Inc platform (Morrisville, North Carolina, US). For metabolite quantification, peaks were quantified using area-under-the-curve. A data normalization step was performed to correct variation resulting from instrument inter-day tuning differences, and values were rescaled to set the medial equal to 1. Missing values were imputed with the minimum.

### Chemical inhibitors, agonists and antagonists

The inhibitor of Syk phosphorylation R406 (Invivogen) and the serine-protease inhibitor 3,4-Dichloroisocoumarin (Sigma Aldrich) were used at 5 μM and 10 μM, respectively. The PPARγ agonist Rosiglitazone (Sigma-Aldrich) and the antagonist GW9662 (Sigma-Aldrich) were used at 10 μM. All drugs were added 1h before BCG stimulation.

### Cytokine stimulation and blocking experiments

Whole blood from healthy donors was pre-incubated with antibodies or a relevant isotype control at 10 μg/mL for 1h (unless otherwise stated), using the following: anti-human CD120a (Clone MABTNFR1-B1, BD Bioscience) antibody, anti-human Interferon alpha/beta receptor 1 antibody (Anifrolumab), anti-human Interferon alpha/beta receptor 2 antibody (PBL, clone MMHAR-2). Whole blood was then diluted 1/3 in TruCulture media and stimulated with BCG or with selected human recombinant cytokines for 22h at 37°C, as follows: IFNα (IntronA; 1000 IU/mL), IFNβ (Betaferon; 1000 IU/mL), IFNγ (Peprotec; 25 ng/mL), IL-1β (Peprotech; 25 ng/mL), TNF (Miltenyi; 10 ng/mL), IL-8 (Biolegend; 25 ng/mL).

### siRNA experiments

Monocytes were seeded at 2×10^5^ cells/200μl in 96-well plates and incubated at 37°C. Cyclophilin B (control) and PPARγ siRNA (SMARTPool, Dharmarcon) were diluted in DOTAP (1,2-dioleoyl-3-trimethylammonium-propane; Roche Applied Sciences). The mix was gently mixed and incubated at room temperature for 15 min. After incubation, the mix was added to cells in culture at a final concentration of 160 nM. Cells were then incubated at 37°C for 24 hours before adding BCG stimulation for 16 hours.

### Flow cytometry – CD36 & PPARγ

Cells were washed in PBS and resuspended in PBS containing 2% fetal calf serum and 2 mM EDTA and stained with the extracellular mix using anti-human CD14-BV421 antibody (clone M5E2, BD Bioscience) and anti-human CD36-BUV605 antibody (clone CD38, BD Bioscience) at 1:200. For PPARγ intracellular staining, a Fixation/Permeabilization Solution Kit (BD Cytofix/Cytoperm) was used according to the manufacturer’s protocol. Briefly, the cells were fixed for 10 min at 4°C with 100 μL of the Fixation/Permeabilization solution and then washed and stained in 100 μL of the BD Perm/Wash Buffer containing anti-human polyclonal PPARγ antibody (PA3-821A, Invitrogen) at 1:25 for 1 hour at 4°C. Goat anti-rabbit IgG-Alexa Fluor 700 was used as a secondary antibody at 1:1000 (ThermoFisher Scientific). Data acquisition was performed on a FACS LSR flow cytometer using FACSDiva software (BD Biosciences, San Jose, CA). FlowJo software (Treestar, Ashland, OR) was used to analyze data.

### Statistical Analysis, Data Visualization, and Software

ANOVA (Kruskal-Wallis) testing was performed for multi-group comparisons and paired t-tests for intra-patient analysis with Qlucore Omics Explorer, v.3.2 (Qlucore) or GraphPAD Prism. To correct for multiple testing we report false discovery rate (FDR)-adjusted ANOVA p values; q values. Dot plot graphs were compiled with GraphPad Prism v.6.0, heatmaps and Principal Component Analysis (PCA) plots with Qlucore. ROC curves were calculated with R (v.3.4.4) and results drawn with graphical package ggplot2 (v.3.1.0). UpSet plots were drawn with UpSetR (v1.3.3), and correlations plots corrplot (v0.84).

Time series analysis on the protein secretion measurements, from 5 different donors, was conducted using a linear mixed-model approach. The time dependency was modelled by incorporating the parameter time as a continuous linear predictor, alongside the other protein predictors, and the donors were modelled as a random effect. The model was implemented using the R package ‘lme” (v1.1-20).

For calculation of cytokine-specific genes score, the genes uniquely induced by each cytokine were identified from our previous study in healthy donors^15^. The first Principal component of the expression matrix was computed using the PCA() function of the FactoMineR package (version 1.42) with the options “scale” set to TRUE, and the option “ncp” set to 1. Position of each sample on the first component was then extracted and used for further analysis (R v.3.6.1).

Pearson correlations were performed in R (v3.6.0) between IL-1Ra protein concentrations quantified by Luminex and the metabolites quantified through mass spectrometry for either TB or LTBI cases. To down-size the number of metabolites analyzed, only metabolites that belong to metabolic groups composed of at least 7 metabolites and that are not labelled as ‘Drug’ or ‘Chemical’ were considered. The metabolite and protein correlation heatmap was plotted with the ggplot2 R package (v.3.3.1).

## Notes

### Competing Interest Statement

MLA is a current employee of Insitro, who had no influence on the study design or reporting. DD has received grant support in the past from Myriad RBM and Roche Genentech but not in the context of this study. The other authors declare no competing interests. 

