## Supplementary figures and images for "Tuberculosis alters immune-metabolic pathways resulting in perturbed IL-1 responses"

### Supplementary figures

#### Supplementary Figure 1

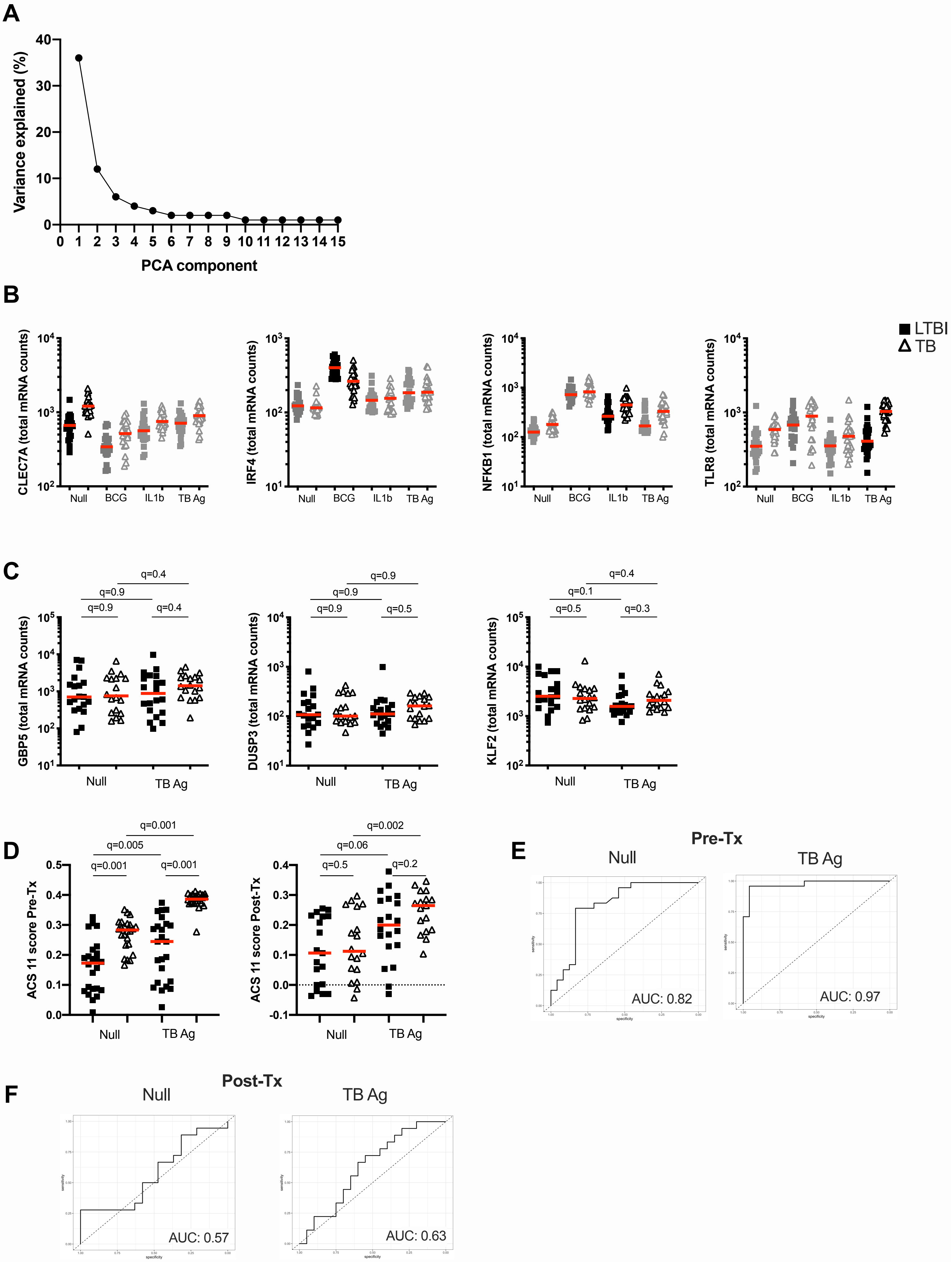

#### Supplementary Figure 2

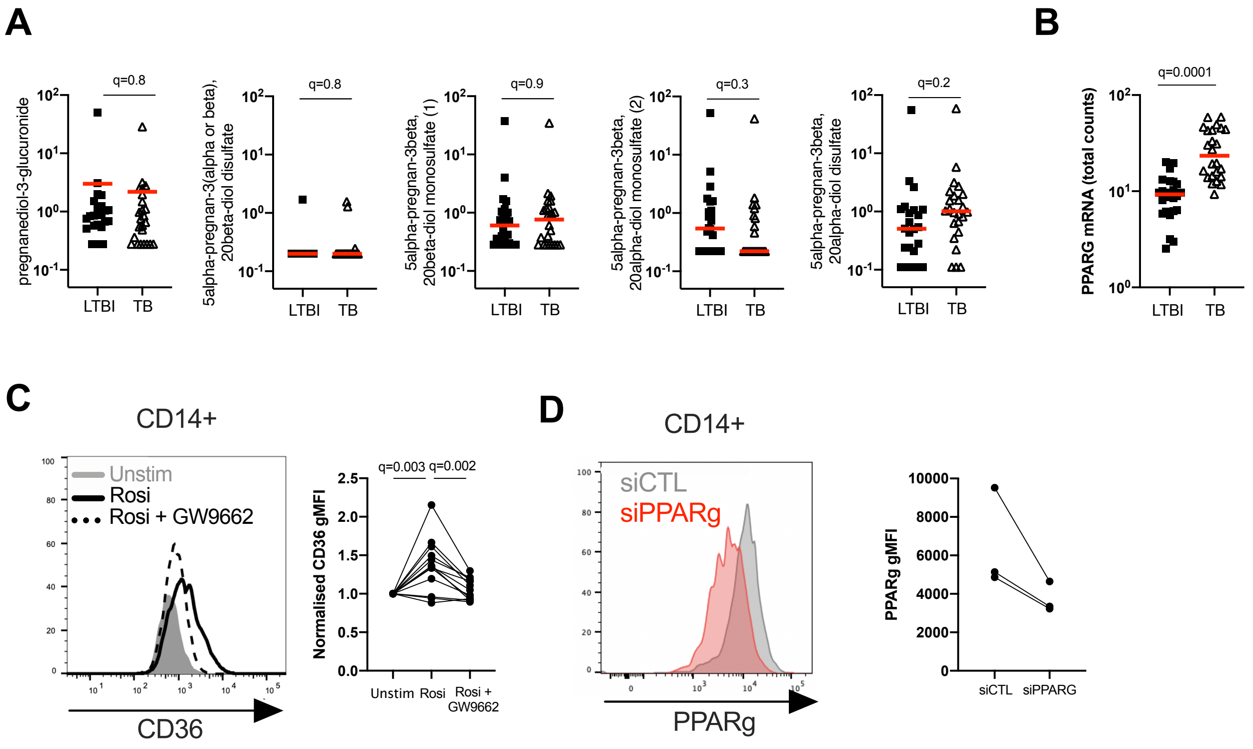
