## Supplementary material for "Tuberculosis alters immune-metabolic pathways resulting in perturbed IL-1 responses": Tables

| | | TB Patients | LTBI controls | p value |
|---|---|---|---|---|
| Age (median years, IQR) | | 33 (25-40) | 33 (24-41) | 0.89 |
| Sex (% female) | | 24 | 24 | >0.99 |
| Ethnicity (% Cape Mixed ancestry) | | 72 | 76 | >0.99 |
| Household TB contacts (%yes) | | 56 | 40 | 0.26 |
| BMI (media, IQR) | | 20 (19-21) | 26 (24-28) | <0.0001 |
| Smoking (n) | Smoker | 14 | 0 | 0.07 |
| | Ex-smoker | 7 | 2 | |
| | Non-smoker | 4 | 10 | |
| Visit 1 (n° donors) | | 24 | 24 | NA |
| Visit 2 (n° donors) | | 18 (after treatment) | 19 | NA |

**Table 1**. Tuberculosis cohort characteristics. TB: tuberculosis; LTBI: Latent tuberculosis infection; IQR: Interquartile range; NA: Non applicable.

| Plus genes |
|---|
| ACTA2 |
| ALDH1A1 |
| ANKRD22 |
| APOL1 |
| APOL6 |
| BATF2 |
| KLF2 |
| CALML4 |
| CASP4 |
| CREB5 |
| CYB561 |
| DEFA1 |
| DUSP3 |
| ETV7 |
| GAS6 |
| GBP2 |
| GBP4 |
| GBP6 |
| HPSE |
| KCNJ15 |
| KREMEN1 |
| LACTB |
| LHFPL2 |
| LOC389386 |
| FER1L3 |
| SCARF1 |
| SEPT4 |
| SMARCD3 |
| TRAFD1 |
| VAMP5 |

**Supplementary Table 1**. Tuberculosis-related genes added to the Nanostring human immunology v2 panel.

| Null (20) | BCG (28) | IL1b (15) | TB Ag (71) | |
|---|---|---|---|---|
| BAX | ADA | C1S | ACTA2 | IL27 |
| CD46 | ALDH1A1 | CR1 | ALAS1 | IL7 |
| CD86 | BATF | CYB561 | ANKRD22 | IRF1 |
| CLEC7A | C7 | DEFB103B | APOL1 | IRF8 |
| CMKLR1 | C8A | DEFB4A | APOL6 | JAK2 |
| CXCL1 | CCL22 | GZMA | BATF2 | KCNJ2 |
| HAVCR2 | CD22 | HLA.DOB | CALML4 | KLRG1 |
| IFNAR2 | CD36 | ICAM2 | CCL24 | LHFPL2 |
| IL10RA | CD99 | KLRK1 | CD274 | LILRB4 |
| IL12RB1 | CLU | NCR1 | CD276 | LOC389386 |
| IL1RL2 | CSF2 | NFKB1 | CD44 | LTB4R2 |
| KIT | CTNNB1 | NOD1 | CD45R0 | MUC1 |
| LILRA2 | CTSG | NT5E | CD74 | PDCD1LG2 |
| MME | IL10 | TNFRSF10C | CD83 | PECAM1 |
| PTGS2 | IL22 | TNFSF12 | CISH | PML |
| RPL19 | IL22RA2 | | CLEC4A | PRDM1 |
| S100A8 | IL6 | | CSF2RB | PSMB10 |
| STAT4 | IRF4 | | CXCL2 | PTAFR |
| STAT6 | JAK1 | | ETV7 | PTPN22 |
| TYK2 | LAIR1 | | FAS | PTPRC |
| | NCF4 | | FCGR1A.B | SMAD3 |
| | PRF1 | | FER1L3 | SOCS1 |
| | PSMB5 | | FN1 | SOCS3 |
| | PSMB7 | | FYN | STAT1 |
| | PSMC2 | | GBP1 | STAT2 |
| | PSMD7 | | GBP4 | STAT5A |
| | TRAF2 | | GFI1 | TAP2 |
| | sCTLA4 | | GPR183 | TGFBI |
| | | | HLA.C | TGFBR2 |
| | | | HLA.DMA | TLR8 |
| | | | HLA.DPB1 | TNFAIP6 |
| | | | HLA.DRA | TNFSF10 |
| | | | IFI16 | TNFSF13B |
| | | | IFITM1 | TRAFD1 |
| | | | IFNB1 | VAMP5 |
| | | | IL21 | |

**Supplementary Table 2**. List of differentially expressed genes between LTBI and TB -only after specific stimulation (t-test with a cut-off value of q<0.001).

| | TB Ag | BCG | |
|---|---|---|---|
| Different | 62 | 40 | |
| No different | 560 | 582 | q=0.03 (*) |

| | TB Ag | IL-1b | |
|---|---|---|---|
| Different | 62 | 16 | |
| No different | 560 | 606 | q=0.0001 (****) |

| | BCG | IL-1b | |
|---|---|---|---|
| Different | 40 | 16 | |
| No different | 582 | 606 | q=0.002 (**) |

**Supplementary Table 3**. Contingency test (Fischer's exact test) for the differentially expressed genes upon immune stimulation (BCG, IL-1β and TB Ag).

| q-value | | | | |
|---|---|---|---|---|
| Transcript | Null | BCG | IL-1b | TB Ag |
| CLL7 | 0.0001 | 0.6145 | 0.4439 | 0.1676 |
| CCL5 | 0.8021 | 0.0006 | 0.6393 | 0.922 |
| NFKB1 | 0.001 | 0.1019 | $2.95 \times 10^{-4}$ | 0.0026 |
| IDO1 | 0.0724 | 0.9277 | 0.443 | $2.24 \times 10^{-5}$ |

**Supplementary Table 4**. Examples of stimuli-specific induced differences between LTBI and TB with q-value specified for each condition.

| BIOCHEMICAL | Pathway | TB | LTBI |
|---|---|---|---|
| 5alpha-pregnan-3beta,20alpha-diol monosulfate (2) | Steroid | 0.55 | -0.06 |
| 5alpha-pregnan-3beta,20beta-diol monosulfate (1) | Steroid | 0.55 | -0.05 |
| 5alpha-pregnan-3beta,20alpha-diol disulfate | Steroid | 0.55 | -0.08 |
| pregnanediol-3-glucuronide | Steroid | 0.51 | -0.04 |
| sphingomyelin (d18:1/20:2, d18:2/20:1, d16:1/22:2)* | Sphingolipid Metabolism | 0.43 | -0.09 |
| 5alpha-pregnan-3(alpha or beta),20beta-diol disulfate | Steroid | 0.40 | 0.11 |
| 4-ethylphenylsulfate | Benzoate Metabolism | 0.39 | -0.02 |
| stearoyl sphingomyelin (d18:1/18:0) | Sphingolipid Metabolism | 0.37 | 0.26 |
| 1-palmitoleoylglycerol (16:1)* | Monoacylglycerol | 0.36 | -0.21 |
| pyrraline | Food Component/Plant | 0.36 | -0.02 |
| tryptophan betaine | Tryptophan Metabolism | 0.35 | -0.33 |
| sphingomyelin (d18:1/22:2, d18:2/22:1, d16:1/24:2)* | Sphingolipid Metabolism | 0.34 | -0.09 |
| 3-hydroxybutyrylcarnitine (2) | Fatty Acid Metabolism(Acyl Carnitine) | 0.33 | 0.20 |
| 1-eicosapentaenoylglycerol (20:5)* | Monoacylglycerol | 0.32 | 0.05 |
| laurate (12:0) | Medium Chain Fatty Acid | 0.32 | 0.12 |
| N-delta-acetylornithine | Urea cycle; Arginine and Proline Metabolism | 0.31 | -0.06 |
| cystine | Methionine, Cysteine, SAM and Taurine Metabolism | 0.30 | -0.10 |
| indole-3-carboxylic acid | Tryptophan Metabolism | 0.29 | 0.14 |
| sphingomyelin (d18:1/18:1, d18:2/18:0) | Sphingolipid Metabolism | 0.29 | 0.25 |
| linoleoyl-docosahexaenoyl-glycerol (18:2/22:6) [1]* | Diacylglycerol | 0.28 | 0.21 |
| 1-linolenoylglycerol (18:3) | Monoacylglycerol | 0.28 | -0.07 |
| 1-arachidonylglycerol (20:4) | Monoacylglycerol | 0.27 | -0.35 |
| 1-docosahexaenoylglycerol (22:6) | Monoacylglycerol | 0.27 | -0.11 |
| betonicine | Food Component/Plant | 0.26 | -0.08 |
| sphingomyelin (d18:0/18:0, d19:0/17:0)* | Sphingolipid Metabolism | 0.26 | 0.48 |
| N-methylproline | Urea cycle; Arginine and Proline Metabolism | 0.24 | -0.07 |
| 4-methylcatechol sulfate | Benzoate Metabolism | 0.24 | 0.03 |
| isobutyrylcarnitine (C4) | Leucine, Isoleucine and Valine Metabolism | 0.24 | 0.16 |
| 1-arachidonoyl-GPA (20:4) | Lysolipid | 0.22 | -0.27 |
| p-cresol sulfate | Phenylalanine and Tyrosine Metabolism | 0.21 | -0.32 |
| gamma-glutamylglutamate | Gamma-glutamyl Amino Acid | 0.20 | -0.30 |
| 2-arachidonoylglycerol (20:4) | Monoacylglycerol | 0.19 | -0.20 |
| 1-palmitoyl-GPA (16:0) | Lysolipid | 0.18 | -0.27 |

| | | | |
|---|---|---|---|
| oleoyl-arachidonoyl-glycerol (18:1/20:4) [1]* | Diacylglycerol | 0.18 | 0.30 |
| stachydrine | Food Component/Plant | 0.17 | -0.06 |
| 1-palmitoyl-2-oleoyl-GPI (16:0/18:1)* | Phospholipid Metabolism | 0.17 | 0.05 |
| 2-palmitoleoyl-GPC (16:1)* | Lysolipid | 0.17 | -0.02 |
| tetradecanedioate | Fatty Acid, Dicarboxylate | 0.17 | 0.33 |
| oleoyl-arachidonoyl-glycerol (18:1/20:4) [2]* | Diacylglycerol | 0.17 | 0.39 |
| 1-palmitoyl-GPG (16:0)* | Lysolipid | 0.16 | 0.15 |
| linoleoyl-arachidonoyl-glycerol (18:2/20:4) [1]* | Diacylglycerol | 0.16 | 0.01 |
| methyl glucopyranoside (alpha + beta) | Food Component/Plant | 0.16 | 0.21 |
| linoleoyl-docosahexaenoyl-glycerol (18:2/22:6) [2]* | Diacylglycerol | 0.16 | 0.23 |
| gluconate | Food Component/Plant | 0.16 | 0.11 |
| 1-palmitoyl-2-linoleoyl-GPI (16:0/18:2) | Phospholipid Metabolism | 0.16 | 0.18 |
| glycocholenate sulfate* | Secondary Bile Acid Metabolism | 0.16 | 0.20 |
| sphingomyelin (d18:2/23:1)* | Sphingolipid Metabolism | 0.16 | 0.24 |
| undecanoate (11:0) | Medium Chain Fatty Acid | 0.15 | -0.01 |
| sphingomyelin (d18:2/24:2)* | Sphingolipid Metabolism | 0.15 | -0.20 |
| indolepropionate | Tryptophan Metabolism | 0.15 | -0.04 |
| 2-palmitoyl-GPC (16:0)* | Lysolipid | 0.15 | -0.21 |
| palmitoleoyl-linoleoyl-glycerol (16:1/18:2) [1]* | Diacylglycerol | 0.14 | 0.18 |
| sphingomyelin (d18:1/17:0, d17:1/18:0, d19:1/16:0) | Sphingolipid Metabolism | 0.14 | 0.38 |
| benzoate | Benzoate Metabolism | 0.14 | 0.21 |
| sphingomyelin (d18:2/18:1)* | Sphingolipid Metabolism | 0.14 | 0.21 |
| 1-myristoylglycerol (14:0) | Monoacylglycerol | 0.13 | -0.13 |
| 3-phenylpropionate (hydrocinnamate) | Phenylalanine and Tyrosine Metabolism | 0.13 | -0.15 |
| sebacate (decanedioate) | Fatty Acid, Dicarboxylate | 0.12 | 0.43 |
| isovalerylcarnitine (C5) | Leucine, Isoleucine and Valine Metabolism | 0.12 | -0.22 |
| choline | Phospholipid Metabolism | 0.12 | 0.31 |
| 1-palmitoyl-GPC (16:0) | Lysolipid | 0.11 | 0.17 |
| gamma-glutamylglutamine | Gamma-glutamyl Amino Acid | 0.11 | -0.18 |
| 1-oleoyl-GPA (18:1) | Lysolipid | 0.11 | -0.24 |
| 4-allylphenol sulfate | Food Component/Plant | 0.11 | -0.26 |
| 1-palmitoleoyl-GPC (16:1)* | Lysolipid | 0.10 | 0.03 |
| gamma-glutamylalanine | Gamma-glutamyl Amino Acid | 0.10 | -0.31 |
| undecanedioate | Fatty Acid, Dicarboxylate | 0.10 | 0.10 |
| docosapentaenoate (n3 DPA; 22:5n3) | Polyunsaturated Fatty Acid (n3 and n6) | 0.10 | 0.07 |
| 2-oxoarginine* | Urea cycle; Arginine and Proline Metabolism | 0.10 | -0.16 |
| isocitrate | TCA Cycle | 0.10 | 0.05 |
| 1-arachidonoyl-GPC (20:4n6)* | Lysolipid | 0.10 | -0.19 |

| | | | |
|---|---|---|---|
| pentadecanoate (15:0) | Long Chain Fatty Acid | 0.10 | 0.19 |
| 3-hydroxyhexanoate | Fatty Acid, Monohydroxy | 0.10 | 0.44 |
| palmitoyl-linolenoyl-glycerol (16:0/18:3) [2]* | Diacylglycerol | 0.10 | 0.32 |
| 1-palmitoylglycerol (16:0) | Monoacylglycerol | 0.10 | -0.32 |
| gentisate | Phenylalanine and Tyrosine Metabolism | 0.09 | 0.11 |
| diacylglycerol (16:1/18:2 [2], 16:0/18:3 [1])* | Diacylglycerol | 0.09 | 0.33 |
| myristoyl dihydrosphingomyelin (d18:0/14:0)* | Sphingolipid Metabolism | 0.09 | 0.23 |
| diacylglycerol (12:0/18:1, 14:0/16:1, 16:0/14:1) [2]* | Diacylglycerol | 0.09 | 0.32 |
| 1-palmitoyl-GPI (16:0) | Lysolipid | 0.08 | -0.10 |
| kynurenine | Tryptophan Metabolism | 0.08 | -0.24 |
| palmitoyl sphingomyelin (d18:1/16:0) | Sphingolipid Metabolism | 0.08 | 0.30 |
| 3-hydroxy-2-ethylpropionate | Leucine, Isoleucine and Valine Metabolism | 0.08 | -0.25 |
| 1-palmitoyl-2-palmitoleoyl-GPC (16:0/16:1)* | Phospholipid Metabolism | 0.08 | 0.25 |
| 4-hydroxyhippurate | Benzoate Metabolism | 0.07 | 0.16 |
| 1-palmitoyl-2-arachidonoyl-GPE (16:0/20:4)* | Phospholipid Metabolism | 0.07 | 0.39 |
| linoleoyl-arachidonoyl-glycerol (18:2/20:4) [2]* | Diacylglycerol | 0.07 | -0.05 |
| alpha-hydroxyisovalerate | Leucine, Isoleucine and Valine Metabolism | 0.07 | -0.05 |
| palmitoyl-myristoyl-glycerol (16:0/14:0) [2] | Diacylglycerol | 0.06 | 0.37 |
| linoleoyl-linolenoyl-glycerol (18:2/18:3) [2]* | Diacylglycerol | 0.06 | 0.06 |
| myristoylcarnitine (C14) | Fatty Acid Metabolism(Acyl Carnitine) | 0.06 | 0.17 |
| isoursodeoxycholate | Secondary Bile Acid Metabolism | 0.05 | -0.02 |
| 3-carboxy-4-methyl-5-propyl-2-furanpropanoate (CMPF) | Fatty Acid, Dicarboxylate | 0.05 | -0.24 |
| 2-stearoyl-GPE (18:0)* | Lysolipid | 0.05 | 0.37 |
| palmitoyl-arachidonoyl-glycerol (16:0/20:4) [2]* | Diacylglycerol | 0.05 | 0.47 |
| myristoleoylcarnitine (C14:1)* | Fatty Acid Metabolism(Acyl Carnitine) | 0.05 | 0.17 |
| linoleoyl-linolenoyl-glycerol (18:2/18:3) [1]* | Diacylglycerol | 0.05 | -0.03 |
| sphingomyelin (d18:1/20:1, d18:2/20:0)* | Sphingolipid Metabolism | 0.04 | -0.04 |
| 1-oleoylglycerol (18:1) | Monoacylglycerol | 0.04 | -0.19 |
| sphingomyelin (d17:1/16:0, d18:1/15:0, d16:1/17:0)* | Sphingolipid Metabolism | 0.04 | 0.33 |
| palmitoleoylcarnitine (C16:1)* | Fatty Acid Metabolism(Acyl Carnitine) | 0.04 | 0.04 |
| sphingomyelin (d18:1/14:0, d16:1/16:0)* | Sphingolipid Metabolism | 0.04 | 0.20 |
| succinylcarnitine (C4-DC) | TCA Cycle | 0.04 | 0.31 |
| glycosyl-N-stearoyl-sphingosine (d18:1/18:0) | Sphingolipid Metabolism | 0.03 | 0.07 |

| | | | |
|---|---|---|---|
| methionine sulfoxide | Methionine, Cysteine, SAM and Taurine Metabolism | 0.03 | -0.19 |
| erythritol | Food Component/Plant | 0.03 | 0.07 |
| 1-palmitoyl-GPE (16:0) | Lysolipid | 0.03 | 0.62 |
| diacylglycerol (14:0/18:1, 16:0/16:1) [2]* | Diacylglycerol | 0.03 | 0.44 |
| 1-stearoyl-2-oleoyl-GPI (18:0/18:1)* | Phospholipid Metabolism | 0.03 | -0.09 |
| docosahexaenoate (DHA; 22:6n3) | Polyunsaturated Fatty Acid (n3 and n6) | 0.02 | 0.12 |
| behenoyl dihydrosphingomyelin (d18:0/22:0)* | Sphingolipid Metabolism | 0.02 | 0.37 |
| sphingomyelin (d18:2/16:0, d18:1/16:1)* | Sphingolipid Metabolism | 0.02 | 0.17 |
| citrulline | Urea cycle; Arginine and Proline Metabolism | 0.02 | -0.03 |
| diacylglycerol (14:0/18:1, 16:0/16:1) [1]* | Diacylglycerol | 0.02 | 0.43 |
| sphingomyelin (d18:2/21:0, d16:2/23:0)* | Sphingolipid Metabolism | 0.02 | 0.29 |
| sphingomyelin (d18:1/19:0, d19:1/18:0)* | Sphingolipid Metabolism | 0.02 | 0.42 |
| 1-linoleoylglycerol (18:2) | Monoacylglycerol | 0.02 | -0.47 |
| 1-palmitoyl-2-oleoyl-GPC (16:0/18:1) | Phospholipid Metabolism | 0.02 | 0.24 |
| dihomo-linoleate (20:2n6) | Polyunsaturated Fatty Acid (n3 and n6) | 0.02 | -0.03 |
| pelargonate (9:0) | Medium Chain Fatty Acid | 0.01 | -0.06 |
| piperine | Food Component/Plant | 0.01 | 0.10 |
| 2-methylbutyrylcarnitine (C5) | Leucine, Isoleucine and Valine Metabolism | 0.01 | 0.09 |
| saccharin | Food Component/Plant | 0.01 | -0.21 |
| p-cresol-glucuronide* | Phenylalanine and Tyrosine Metabolism | 0.01 | 0.15 |
| betaine | Glycine, Serine and Threonine Metabolism | 0.01 | -0.19 |
| 1-palmitoleoyl-2-linoleoyl-GPC (16:1/18:2)* | Phospholipid Metabolism | 0.01 | -0.02 |
| 3-indoxyl sulfate | Tryptophan Metabolism | 0.00 | -0.18 |
| oleoyl-oleoyl-glycerol (18:1/18:1) [2]* | Diacylglycerol | 0.00 | 0.50 |
| glycolithocholate sulfate* | Secondary Bile Acid Metabolism | 0.00 | -0.01 |
| glycoursodeoxycholate | Secondary Bile Acid Metabolism | 0.00 | 0.05 |
| 3-methoxytyrosine | Phenylalanine and Tyrosine Metabolism | 0.00 | -0.08 |
| 1-linoleoyl-GPA (18:2)* | Lysolipid | 0.00 | -0.31 |
| 2-palmitoylglycerol (16:0) | Monoacylglycerol | 0.00 | -0.21 |
| 1-stearoyl-GPE (18:0) | Lysolipid | 0.00 | 0.61 |
| linoleate (18:2n6) | Polyunsaturated Fatty Acid (n3 and n6) | 0.00 | 0.03 |
| proline | Urea cycle; Arginine and Proline Metabolism | 0.00 | -0.01 |
| 1-palmitoyl-2-linoleoyl-GPE (16:0/18:2) | Phospholipid Metabolism | 0.00 | 0.55 |
| picolinate | Tryptophan Metabolism | 0.00 | 0.28 |
| N-acetylvaline | Leucine, Isoleucine and Valine Metabolism | -0.61 | 0.16 |

| | | | |
|---|---|---|---|
| N-acetylleucine | Leucine, Isoleucine and Valine Metabolism | -0.54 | -0.04 |
| N-acetylglutamine | Glutamate Metabolism | -0.52 | -0.30 |
| gamma-glutamylthreonine | Gamma-glutamyl Amino Acid | -0.45 | -0.15 |
| 3-(4-hydroxyphenyl)lactate | Phenylalanine and Tyrosine Metabolism | -0.45 | -0.22 |
| 2-hydroxyoctanoate | Fatty Acid, Monohydroxy | -0.44 | -0.16 |
| indolelactate | Tryptophan Metabolism | -0.42 | -0.20 |
| gamma-glutamylleucine | Gamma-glutamyl Amino Acid | -0.42 | -0.13 |
| gamma-glutamyltyrosine | Gamma-glutamyl Amino Acid | -0.42 | -0.20 |
| 2-keto-3-deoxy-gluconate | Food Component/Plant | -0.41 | 0.07 |
| N-acetylisoleucine | Leucine, Isoleucine and Valine Metabolism | -0.41 | -0.17 |
| gamma-glutamyl-epsilon-lysine | Gamma-glutamyl Amino Acid | -0.40 | -0.02 |
| gamma-glutamylvaline | Gamma-glutamyl Amino Acid | -0.39 | 0.13 |
| 13-HODE + 9-HODE | Fatty Acid, Monohydroxy | -0.39 | 0.08 |
| 1-oleoyl-GPS (18:1) | Lysolipid | -0.39 | -0.04 |
| cysteine | Methionine, Cysteine, SAM and Taurine Metabolism | -0.38 | -0.03 |
| imidazole lactate | Histidine Metabolism | -0.38 | -0.09 |
| alliin | Food Component/Plant | -0.38 | -0.09 |
| valine | Leucine, Isoleucine and Valine Metabolism | -0.37 | 0.13 |
| tryptophan | Tryptophan Metabolism | -0.36 | -0.15 |
| N-acetylarginine | Urea cycle; Arginine and Proline Metabolism | -0.36 | 0.07 |
| thymol sulfate | Food Component/Plant | -0.36 | 0.26 |
| 5-acetylamino-6-formylamino-3-methyluracil | Xanthine Metabolism | -0.36 | -0.19 |
| trans-urocanate | Histidine Metabolism | -0.35 | -0.07 |
| pyroglutamine* | Glutamate Metabolism | -0.35 | -0.23 |
| 3-methylxanthine | Xanthine Metabolism | -0.35 | 0.22 |
| trans-4-hydroxyproline | Urea cycle; Arginine and Proline Metabolism | -0.35 | 0.24 |
| N-acetylglutamate | Glutamate Metabolism | -0.35 | 0.45 |
| N-acetylphenylalanine | Phenylalanine and Tyrosine Metabolism | -0.34 | -0.44 |
| glycerophosphorylcholine (GPC) | Phospholipid Metabolism | -0.34 | 0.18 |
| glycerophosphoethanolamine | Phospholipid Metabolism | -0.34 | 0.27 |
| N-formylmethionine | Methionine, Cysteine, SAM and Taurine Metabolism | -0.34 | 0.17 |
| gamma-glutamylisoleucine* | Gamma-glutamyl Amino Acid | -0.34 | 0.00 |
| N-acetylhistidine | Histidine Metabolism | -0.33 | -0.05 |
| theobromine | Xanthine Metabolism | -0.33 | -0.08 |
| kynurenate | Tryptophan Metabolism | -0.32 | -0.19 |
| glutamate | Glutamate Metabolism | -0.32 | -0.03 |
| threonine | Glycine, Serine and Threonine Metabolism | -0.32 | -0.13 |

| | | | |
|---|---|---|---|
| 5alpha-androstan-3beta,17beta-diol monosulfate (2) | Steroid | -0.32 | -0.33 |
| 1-linoleoyl-GPG (18:2)* | Lysolipid | -0.32 | -0.24 |
| glutamine | Glutamate Metabolism | -0.31 | -0.02 |
| 1,7-dimethylurate | Xanthine Metabolism | -0.31 | 0.04 |
| choline phosphate | Phospholipid Metabolism | -0.31 | 0.04 |
| malate | TCA Cycle | -0.31 | 0.10 |
| 2-hydroxyglutarate | Fatty Acid, Dicarboxylate | -0.31 | 0.13 |
| cortisol | Steroid | -0.31 | 0.38 |
| paraxanthine | Xanthine Metabolism | -0.30 | 0.06 |
| homoarginine | Urea cycle; Arginine and Proline Metabolism | -0.30 | -0.20 |
| 5alpha-androstan-3alpha,17beta-diol monosulfate (1) | Steroid | -0.30 | -0.27 |
| N-acetylserine | Glycine, Serine and Threonine Metabolism | -0.30 | 0.32 |
| N-acetyltyrosine | Phenylalanine and Tyrosine Metabolism | -0.30 | -0.17 |
| gamma-glutamyl-alpha-lysine | Gamma-glutamyl Amino Acid | -0.30 | -0.02 |
| dimethylarginine (SDMA + ADMA) | Urea cycle; Arginine and Proline Metabolism | -0.29 | 0.15 |
| 1,3,7-trimethylurate | Xanthine Metabolism | -0.29 | -0.15 |
| 1-linoleoyl-GPE (18:2)* | Lysolipid | -0.29 | -0.06 |
| 1,2-dilinoleoyl-GPC (18:2/18:2) | Phospholipid Metabolism | -0.29 | -0.16 |
| 1-methylhistidine | Histidine Metabolism | -0.29 | 0.04 |
| 1-stearoyl-2-linoleoyl-GPC (18:0/18:2)* | Phospholipid Metabolism | -0.29 | 0.03 |
| N-behenoyl-sphingadienine (d18:2/22:0)* | Sphingolipid Metabolism | -0.29 | 0.18 |
| 1-methylxanthine | Xanthine Metabolism | -0.29 | 0.19 |
| S-adenosylhomocysteine (SAH) | Methionine, Cysteine, SAM and Taurine Metabolism | -0.29 | -0.02 |
| 2-methylcitrate/homocitrate | TCA Cycle | -0.28 | -0.10 |
| alpha-hydroxyisocaproate | Leucine, Isoleucine and Valine Metabolism | -0.28 | 0.03 |
| lactosyl-N-nervonoyl-sphingosine (d18:1/24:1)* | Sphingolipid Metabolism | -0.28 | -0.17 |
| urea | Urea cycle; Arginine and Proline Metabolism | -0.28 | 0.40 |
| gamma-glutamylhistidine | Gamma-glutamyl Amino Acid | -0.28 | -0.21 |
| 3-hydroxyhippurate | Benzoate Metabolism | -0.27 | -0.09 |
| 3-methyl-2-oxovalerate | Leucine, Isoleucine and Valine Metabolism | -0.27 | 0.14 |
| adipate | Fatty Acid, Dicarboxylate | -0.27 | 0.10 |
| propyl 4-hydroxybenzoate sulfate | Benzoate Metabolism | -0.27 | 0.03 |
| glycosyl-N-palmitoyl-sphingosine (d18:1/16:0) | Sphingolipid Metabolism | -0.27 | 0.33 |
| citrate | TCA Cycle | -0.27 | -0.23 |
| lignoceroyl sphingomyelin (d18:1/24:0) | Sphingolipid Metabolism | -0.26 | 0.06 |
| androstenediol (3alpha, 17alpha) monsulfate (2) | Steroid | -0.26 | -0.20 |
| behenoyl sphingomyelin (d18:1/22:0)* | Sphingolipid Metabolism | -0.26 | 0.28 |

| | | | |
|---|---|---|---|
| 4-hydroxyphenylpyruvate | Phenylalanine and Tyrosine Metabolism | -0.26 | 0.39 |
| succinate | TCA Cycle | -0.26 | 0.28 |
| 1-arachidonoyl-GPE (20:4n6)* | Lysolipid | -0.26 | 0.08 |
| S-methylcysteine | Methionine, Cysteine, SAM and Taurine Metabolism | -0.26 | -0.35 |
| N-acetyl-1-methylhistidine* | Histidine Metabolism | -0.26 | 0.12 |
| octadecanedioate | Fatty Acid, Dicarboxylate | -0.25 | 0.42 |
| fumarate | TCA Cycle | -0.25 | 0.08 |
| 2-hydroxydecanoate | Fatty Acid, Monohydroxy | -0.25 | -0.11 |
| sarcosine | Glycine, Serine and Threonine Metabolism | -0.25 | 0.01 |
| 5-bromotryptophan | Phenylalanine and Tyrosine Metabolism | -0.25 | -0.34 |
| hypotaurine | Methionine, Cysteine, SAM and Taurine Metabolism | -0.25 | 0.04 |
| 1-arachidonoyl-GPI (20:4)* | Lysolipid | -0.25 | -0.11 |
| androstenediol (3beta,17beta) disulfate (2) | Steroid | -0.25 | -0.20 |
| taurine | Methionine, Cysteine, SAM and Taurine Metabolism | -0.24 | -0.07 |
| phenylalanine | Phenylalanine and Tyrosine Metabolism | -0.24 | 0.01 |
| 3-methyl-2-oxobutyrate | Leucine, Isoleucine and Valine Metabolism | -0.24 | 0.15 |
| cerotoylcarnitine (C26)* | Fatty Acid Metabolism(Acyl Carnitine) | -0.24 | 0.17 |
| 5alpha-androstan-3beta,17alpha-diol disulfate | Steroid | -0.24 | -0.49 |
| 16-hydroxypalmitate | Fatty Acid, Monohydroxy | -0.24 | 0.34 |
| leucine | Leucine, Isoleucine and Valine Metabolism | -0.24 | 0.11 |
| tyrosine | Phenylalanine and Tyrosine Metabolism | -0.23 | 0.09 |
| 8-hydroxyoctanoate | Fatty Acid, Monohydroxy | -0.23 | 0.20 |
| 1-linoleoyl-GPI (18:2)* | Lysolipid | -0.23 | 0.03 |
| maleate | Fatty Acid, Dicarboxylate | -0.23 | 0.58 |
| 3-hydroxylaurate | Fatty Acid, Monohydroxy | -0.23 | 0.59 |
| glycohyocholate | Secondary Bile Acid Metabolism | -0.23 | 0.11 |
| trimethylamine N-oxide | Phospholipid Metabolism | -0.23 | 0.29 |
| stearoylcarnitine (C18) | Fatty Acid Metabolism(Acyl Carnitine) | -0.23 | 0.19 |
| gamma-glutamylphenylalanine | Gamma-glutamyl Amino Acid | -0.22 | -0.18 |
| azelate (nonanedioate) | Fatty Acid, Dicarboxylate | -0.22 | -0.20 |
| isovalerylglycine | Leucine, Isoleucine and Valine Metabolism | -0.22 | -0.07 |
| catechol sulfate | Benzoate Metabolism | -0.22 | -0.07 |
| theophylline | Xanthine Metabolism | -0.22 | -0.03 |
| 5alpha-androstan-3alpha,17beta-diol disulfate | Steroid | -0.22 | -0.01 |
| nervonate (24:1n9)* | Long Chain Fatty Acid | -0.22 | 0.18 |

| | | | |
|---|---|---|---|
| 1-(1-enyl-palmitoyl)-2-arachidonoyl-GPC (P-16:0/20:4)* | Plasmalogen | -0.22 | -0.10 |
| eicosanodioate | Fatty Acid, Dicarboxylate | -0.22 | -0.25 |
| o-cresol sulfate | Phenylalanine and Tyrosine Metabolism | -0.21 | 0.06 |
| C-glycosyltryptophan | Tryptophan Metabolism | -0.21 | -0.19 |
| behenate (22:0)* | Long Chain Fatty Acid | -0.21 | -0.25 |
| sphinganine-1-phosphate | Sphingolipid Metabolism | -0.21 | -0.09 |
| dihomo-linoleoylcarnitine (C20:2)* | Fatty Acid Metabolism(Acyl Carnitine) | -0.21 | 0.00 |
| S-methylcysteine sulfoxide | Methionine, Cysteine, SAM and Taurine Metabolism | -0.21 | -0.09 |
| etiocholanolone glucuronide | Steroid | -0.21 | 0.07 |
| quinate | Food Component/Plant | -0.20 | -0.11 |
| N-acetylmethionine | Methionine, Cysteine, SAM and Taurine Metabolism | -0.20 | 0.01 |
| indoleacetate | Tryptophan Metabolism | -0.20 | -0.01 |
| formiminoglutamate | Histidine Metabolism | -0.20 | 0.33 |
| caffeine | Xanthine Metabolism | -0.20 | -0.01 |
| sphingomyelin (d18:2/14:0, d18:1/14:1)* | Sphingolipid Metabolism | -0.20 | -0.07 |
| docosadioate | Fatty Acid, Dicarboxylate | -0.20 | -0.13 |
| dimethylglycine | Glycine, Serine and Threonine Metabolism | -0.20 | 0.18 |
| N-acetyltryptophan | Tryptophan Metabolism | -0.20 | -0.19 |
| 7-methylurate | Xanthine Metabolism | -0.20 | 0.18 |
| docosapentaenoate (n6 DPA; 22:5n6) | Polyunsaturated Fatty Acid (n3 and n6) | -0.20 | 0.04 |
| hyocholate | Secondary Bile Acid Metabolism | -0.20 | 0.12 |
| 3-methyl catechol sulfate (1) | Benzoate Metabolism | -0.19 | 0.03 |
| androstenediol (3beta,17beta) disulfate (1) | Steroid | -0.19 | -0.28 |
| 1-stearoyl-2-linoleoyl-GPI (18:0/18:2) | Phospholipid Metabolism | -0.19 | 0.11 |
| 1-(1-enyl-palmitoyl)-2-linoleoyl-GPE (P-16:0/18:2)* | Plasmalogen | -0.19 | 0.33 |
| O-methylcatechol sulfate | Benzoate Metabolism | -0.19 | 0.08 |
| N-acetyltaurine | Methionine, Cysteine, SAM and Taurine Metabolism | -0.19 | -0.36 |
| androsterone sulfate | Steroid | -0.19 | -0.36 |
| argininosuccinate | Urea cycle; Arginine and Proline Metabolism | -0.19 | 0.43 |
| pregnen-diol disulfate* | Steroid | -0.19 | -0.22 |
| 1-oleoyl-GPE (18:1) | Lysolipid | -0.18 | 0.15 |
| ximenoylcarnitine (C26:1)* | Fatty Acid Metabolism(Acyl Carnitine) | -0.18 | 0.04 |
| docosapentaenoylcarnitine (C22:5n3)* | Fatty Acid Metabolism(Acyl Carnitine) | -0.18 | 0.00 |
| myristoleate (14:1n5) | Long Chain Fatty Acid | -0.18 | 0.30 |
| 1-linoleoyl-2-linolenoyl-GPC (18:2/18:3)* | Phospholipid Metabolism | -0.18 | 0.07 |
| androstenediol (3beta,17beta) monosulfate (2) | Steroid | -0.18 | -0.38 |

| | | | |
|---|---|---|---|
| 1-(1-enyl-palmitoyl)-2-palmitoyl-GPC (P-16:0/16:0)* | Plasmalogen | -0.18 | 0.04 |
| eugenol sulfate | Food Component/Plant | -0.18 | 0.39 |
| 1-linoleoyl-GPC (18:2) | Lysolipid | -0.18 | -0.28 |
| cysteine s-sulfate | Methionine, Cysteine, SAM and Taurine Metabolism | -0.18 | -0.07 |
| methyl-4-hydroxybenzoate sulfate | Benzoate Metabolism | -0.18 | -0.09 |
| N-alpha-acetylornithine | Urea cycle; Arginine and Proline Metabolism | -0.18 | -0.02 |
| tricosanoyl sphingomyelin (d18:1/23:0)* | Sphingolipid Metabolism | -0.17 | 0.34 |
| arachidoylcarnitine (C20)* | Fatty Acid Metabolism(Acyl Carnitine) | -0.17 | 0.19 |
| linoleoylcarnitine (C18:2)* | Fatty Acid Metabolism(Acyl Carnitine) | -0.17 | -0.11 |
| 1-linoleoyl-2-arachidonoyl-GPC (18:2/20:4n6)* | Phospholipid Metabolism | -0.17 | 0.01 |
| lignoceroylcarnitine (C24)* | Fatty Acid Metabolism(Acyl Carnitine) | -0.17 | -0.04 |
| 1,3-dimethylurate | Xanthine Metabolism | -0.17 | -0.09 |
| palmitoyl-palmitoyl-glycerol (16:0/16:0) [1]* | Diacylglycerol | -0.17 | 0.45 |
| androstenediol (3beta,17beta) monosulfate (1) | Steroid | -0.17 | -0.14 |
| pregn steroid monosulfate* | Steroid | -0.17 | -0.28 |
| 5alpha-androstan-3beta,17beta-diol disulfate | Steroid | -0.17 | -0.34 |
| N-acetylthreonine | Glycine, Serine and Threonine Metabolism | -0.17 | 0.11 |
| 1-methylimidazoleacetate | Histidine Metabolism | -0.17 | 0.06 |
| stearate (18:0) | Long Chain Fatty Acid | -0.16 | 0.14 |
| 2-oleoylglycerol (18:1) | Monoacylglycerol | -0.16 | -0.08 |
| 3-hydroxydecanoate | Fatty Acid, Monohydroxy | -0.16 | 0.69 |
| cortisone | Steroid | -0.16 | -0.22 |
| epiandrosterone sulfate | Steroid | -0.16 | -0.36 |
| 1-palmitoyl-2-stearoyl-GPC (16:0/18:0) | Phospholipid Metabolism | -0.15 | 0.41 |
| sphingomyelin (d18:1/24:1, d18:2/24:0)* | Sphingolipid Metabolism | -0.15 | -0.09 |
| caprylate (8:0) | Medium Chain Fatty Acid | -0.15 | -0.36 |
| thyroxine | Phenylalanine and Tyrosine Metabolism | -0.15 | -0.17 |
| 5-acetylamino-6-amino-3-methyluracil | Xanthine Metabolism | -0.15 | 0.02 |
| stearidonate (18:4n3) | Polyunsaturated Fatty Acid (n3 and n6) | -0.15 | 0.25 |
| dopamine 3-O-sulfate | Phenylalanine and Tyrosine Metabolism | -0.15 | -0.27 |
| methionine | Methionine, Cysteine, SAM and Taurine Metabolism | -0.15 | -0.24 |
| sphingomyelin (d18:1/22:1, d18:2/22:0, d16:1/24:1)* | Sphingolipid Metabolism | -0.15 | 0.06 |
| erucate (22:1n9) | Long Chain Fatty Acid | -0.15 | 0.01 |
| N-(2-furoyl)glycine | Food Component/Plant | -0.15 | -0.23 |
| beta-citrylglutamate | Glutamate Metabolism | -0.15 | 0.05 |

| | | | |
|---|---|---|---|
| 1-stearoyl-2-arachidonoyl-GPE (18:0/20:4) | Phospholipid Metabolism | -0.15 | 0.35 |
| 4-methyl-2-oxopentanoate | Leucine, Isoleucine and Valine Metabolism | -0.15 | 0.19 |
| N-formylphenylalanine | Phenylalanine and Tyrosine Metabolism | -0.15 | -0.02 |
| 1-palmitoyl-2-linoleoyl-GPC (16:0/18:2) | Phospholipid Metabolism | -0.14 | 0.26 |
| andro steroid monosulfate (1)* | Steroid | -0.14 | -0.30 |
| S-1-pyrroline-5-carboxylate | Glutamate Metabolism | -0.14 | -0.07 |
| 5-dodecenoate (12:1n7) | Medium Chain Fatty Acid | -0.14 | 0.34 |
| serine | Glycine, Serine and Threonine Metabolism | -0.14 | -0.45 |
| adrenoylcarnitine (C22:4)* | Fatty Acid Metabolism(Acyl Carnitine) | -0.14 | -0.07 |
| sphingomyelin (d18:1/25:0, d19:0/24:1, d20:1/23:0, d19:1/24:0)* | Sphingolipid Metabolism | -0.14 | 0.19 |
| 1-oleoyl-GPI (18:1)* | Lysolipid | -0.14 | -0.12 |
| dihomo-linolenate (20:3n3 or n6) | Polyunsaturated Fatty Acid (n3 and n6) | -0.14 | -0.19 |
| 2-hydroxy-3-methylvalerate | Leucine, Isoleucine and Valine Metabolism | -0.14 | -0.02 |
| ornithine | Urea cycle; Arginine and Proline Metabolism | -0.14 | -0.14 |
| imidazole propionate | Histidine Metabolism | -0.13 | 0.23 |
| eicosenoylcarnitine (C20:1)* | Fatty Acid Metabolism(Acyl Carnitine) | -0.13 | 0.11 |
| 1-stearoyl-GPI (18:0) | Lysolipid | -0.13 | -0.20 |
| docosadienoate (22:2n6) | Polyunsaturated Fatty Acid (n3 and n6) | -0.13 | 0.10 |
| 1-(1-enyl-stearoyl)-2-arachidonoyl-GPE (P-18:0/20:4)* | Plasmalogen | -0.13 | 0.44 |
| acetylcarnitine (C2) | Fatty Acid Metabolism(Acyl Carnitine) | -0.13 | 0.45 |
| arachidate (20:0) | Long Chain Fatty Acid | -0.13 | -0.08 |
| 2-hydroxyhippurate (salicylurate) | Benzoate Metabolism | -0.13 | 0.29 |
| glycine | Glycine, Serine and Threonine Metabolism | -0.13 | -0.10 |
| 5-hydroxymethyl-2-furoic acid | Phenylalanine and Tyrosine Metabolism | -0.13 | -0.18 |
| 2-hydroxystearate | Fatty Acid, Monohydroxy | -0.13 | -0.03 |
| gamma-glutamyltryptophan | Gamma-glutamyl Amino Acid | -0.13 | -0.21 |
| 1-(1-enyl-palmitoyl)-2-linoleoyl-GPC (P-16:0/18:2)* | Plasmalogen | -0.12 | -0.09 |
| sphingosine 1-phosphate | Sphingolipid Metabolism | -0.12 | -0.29 |
| palmitoyl-oleoyl-glycerol (16:0/18:1) [1]* | Diacylglycerol | -0.12 | 0.63 |
| 1-(1-enyl-palmitoyl)-2-oleoyl-GPC (P-16:0/18:1)* | Plasmalogen | -0.12 | -0.05 |
| histidine | Histidine Metabolism | -0.12 | 0.00 |
| sphingomyelin (d18:2/23:0, d18:1/23:1, d17:1/24:1)* | Sphingolipid Metabolism | -0.12 | 0.10 |
| glycodeoxycholate | Secondary Bile Acid Metabolism | -0.12 | 0.25 |
| phenol sulfate | Phenylalanine and Tyrosine Metabolism | -0.12 | -0.37 |

| | | | |
|---|---|---|---|
| hippurate | Benzoate Metabolism | -0.12 | 0.20 |
| octanoylcarnitine (C8) | Fatty Acid Metabolism(Acyl Carnitine) | -0.12 | 0.44 |
| cinnamoylglycine | Food Component/Plant | -0.12 | -0.10 |
| margaroylcarnitine* | Fatty Acid Metabolism(Acyl Carnitine) | -0.12 | 0.25 |
| 1,2-dipalmitoyl-GPC (16:0/16:0) | Phospholipid Metabolism | -0.12 | 0.40 |
| linolenate [alpha or gamma; (18:3n3 or 6)] | Polyunsaturated Fatty Acid (n3 and n6) | -0.11 | 0.09 |
| 3-hydroxystearate | Fatty Acid, Monohydroxy | -0.11 | 0.19 |
| androstenediol (3alpha, 17alpha) monsulfate (3) | Steroid | -0.11 | -0.32 |
| alpha-ketoglutarate | TCA Cycle | -0.11 | 0.22 |
| argininate* | Urea cycle; Arginine and Proline Metabolism | -0.11 | 0.36 |
| eicosenoate (20:1) | Long Chain Fatty Acid | -0.11 | 0.11 |
| 1-stearoyl-2-arachidonoyl-GPC (18:0/20:4) | Phospholipid Metabolism | -0.11 | -0.19 |
| oleate/vaccenate (18:1) | Long Chain Fatty Acid | -0.11 | 0.22 |
| phenyllactate (PLA) | Phenylalanine and Tyrosine Metabolism | -0.11 | -0.07 |
| myristate (14:0) | Long Chain Fatty Acid | -0.11 | 0.20 |
| arachidonoylcarnitine (C20:4) | Fatty Acid Metabolism(Acyl Carnitine) | -0.11 | -0.07 |
| oleoylcarnitine (C18:1) | Fatty Acid Metabolism(Acyl Carnitine) | -0.11 | -0.14 |
| 1-(1-enyl-palmitoyl)-2-arachidonoyl-GPE (P-16:0/20:4)* | Plasmalogen | -0.11 | 0.21 |
| N-palmitoyl-sphinganine (d18:0/16:0) | Sphingolipid Metabolism | -0.11 | 0.45 |
| sphingosine | Sphingolipid Metabolism | -0.11 | -0.22 |
| 1-linolenoyl-GPC (18:3)* | Lysolipid | -0.10 | -0.03 |
| N-acetylglycine | Glycine, Serine and Threonine Metabolism | -0.10 | 0.01 |
| hexadecanedioate | Fatty Acid, Dicarboxylate | -0.10 | 0.51 |
| methylsuccinate | Leucine, Isoleucine and Valine Metabolism | -0.10 | 0.25 |
| N-stearoyl-sphingosine (d18:1/18:0)* | Sphingolipid Metabolism | -0.10 | 0.62 |
| pro-hydroxy-pro | Urea cycle; Arginine and Proline Metabolism | -0.10 | 0.19 |
| taurodeoxycholate | Secondary Bile Acid Metabolism | -0.10 | 0.14 |
| 1-stearoyl-2-arachidonoyl-GPI (18:0/20:4) | Phospholipid Metabolism | -0.10 | -0.20 |
| 1-palmitoyl-2-arachidonoyl-GPI (16:0/20:4)* | Phospholipid Metabolism | -0.10 | 0.00 |
| palmitoyl-oleoyl-glycerol (16:0/18:1) [2]* | Diacylglycerol | -0.10 | 0.64 |
| methionine sulfone | Methionine, Cysteine, SAM and Taurine Metabolism | -0.10 | -0.06 |
| phosphoethanolamine | Phospholipid Metabolism | -0.10 | 0.23 |
| 1-stearoyl-GPS (18:0)* | Lysolipid | -0.10 | -0.14 |
| 1-dihomo-linolenylglycerol (20:3) | Monoacylglycerol | -0.09 | -0.49 |
| palmitoyl dihydrosphingomyelin (d18:0/16:0)* | Sphingolipid Metabolism | -0.09 | 0.31 |

| | | | |
|---|---|---|---|
| 2-ethylphenylsulfate | Benzoate Metabolism | -0.09 | 0.09 |
| cis-aconitate | TCA Cycle | -0.09 | -0.15 |
| palmitoleate (16:1n7) | Long Chain Fatty Acid | -0.09 | 0.19 |
| dihomo-linolenoylcarnitine (20:3n3 or 6)* | Fatty Acid Metabolism(Acyl Carnitine) | -0.09 | -0.11 |
| decanoylcarnitine (C10) | Fatty Acid Metabolism(Acyl Carnitine) | -0.09 | 0.34 |
| 3-methoxycatechol sulfate (1) | Benzoate Metabolism | -0.09 | 0.21 |
| ergothioneine | Food Component/Plant | -0.09 | -0.12 |
| heptanoate (7:0) | Medium Chain Fatty Acid | -0.09 | 0.27 |
| N-palmitoyl-sphingosine (d18:1/16:0) | Sphingolipid Metabolism | -0.09 | 0.41 |
| 10-nonadecenoate (19:1n9) | Long Chain Fatty Acid | -0.09 | 0.24 |
| taurocholenate sulfate | Secondary Bile Acid Metabolism | -0.09 | 0.14 |
| stearoyl-arachidonoyl-glycerol (18:0/20:4) [2]* | Diacylglycerol | -0.09 | 0.48 |
| 4-vinylphenol sulfate | Benzoate Metabolism | -0.08 | -0.06 |
| 1-stearoyl-GPC (18:0) | Lysolipid | -0.08 | 0.06 |
| 1-stearoyl-2-oleoyl-GPC (18:0/18:1) | Phospholipid Metabolism | -0.08 | 0.26 |
| glycodeoxycholate sulfate | Secondary Bile Acid Metabolism | -0.08 | 0.27 |
| 1-stearoyl-2-oleoyl-GPE (18:0/18:1) | Phospholipid Metabolism | -0.08 | 0.37 |
| 1-(1-enyl-palmitoyl)-2-oleoyl-GPE (P-16:0/18:1)* | Plasmalogen | -0.08 | 0.20 |
| nonadecanoate (19:0) | Long Chain Fatty Acid | -0.08 | 0.00 |
| gamma-glutamylmethionine | Gamma-glutamyl Amino Acid | -0.08 | -0.26 |
| oleoyl-oleoyl-glycerol (18:1/18:1) [1]* | Diacylglycerol | -0.08 | 0.44 |
| arachidonate (20:4n6) | Polyunsaturated Fatty Acid (n3 and n6) | -0.08 | -0.11 |
| N-acetylproline | Urea cycle; Arginine and Proline Metabolism | -0.07 | 0.00 |
| palmitoyl-palmitoyl-glycerol (16:0/16:0) [2]* | Diacylglycerol | -0.07 | 0.54 |
| 2-hydroxybutyrate/2-hydroxyisobutyrate | Methionine, Cysteine, SAM and Taurine Metabolism | -0.07 | 0.40 |
| 14-HDoHE/17-HDoHE | Fatty Acid, Monohydroxy | -0.07 | 0.25 |
| palmitoylcarnitine (C16) | Fatty Acid Metabolism(Acyl Carnitine) | -0.07 | 0.05 |
| 16a-hydroxy DHEA 3-sulfate | Steroid | -0.07 | -0.22 |
| 2-hydroxypalmitate | Fatty Acid, Monohydroxy | -0.07 | 0.11 |
| glycerophosphoinositol* | Phospholipid Metabolism | -0.07 | 0.00 |
| 3-hydroxyisobutyrate | Leucine, Isoleucine and Valine Metabolism | -0.07 | -0.20 |
| laurylcarnitine (C12) | Fatty Acid Metabolism(Acyl Carnitine) | -0.06 | 0.31 |
| 2-hydroxylaurate | Fatty Acid, Monohydroxy | -0.06 | -0.06 |
| 1-(1-enyl-palmitoyl)-2-palmitoleoyl-GPC (P-16:0/16:1)* | Plasmalogen | -0.06 | 0.01 |
| linoleoyl-linoleoyl-glycerol (18:2/18:2) [1]* | Diacylglycerol | -0.06 | 0.03 |
| taurolithocholate 3-sulfate | Secondary Bile Acid Metabolism | -0.06 | 0.05 |
| 2-hydroxyphenylacetate | Phenylalanine and Tyrosine Metabolism | -0.06 | 0.38 |

| | | | |
|---|---|---|---|
| cis-4-decenoylcarnitine (C10:1) | Fatty Acid Metabolism(Acyl Carnitine) | -0.06 | 0.22 |
| palmitoyl-linoleoyl-glycerol (16:0/18:2) [2]* | Diacylglycerol | -0.06 | 0.53 |
| sphingomyelin (d18:1/20:0, d16:1/22:0)* | Sphingolipid Metabolism | -0.06 | 0.33 |
| lactosyl-N-palmitoyl-sphingosine (d18:1/16:0) | Sphingolipid Metabolism | -0.06 | 0.05 |
| dehydroisoandrosterone sulfate (DHEA-S) | Steroid | -0.06 | -0.29 |
| ethylmalonate | Leucine, Isoleucine and Valine Metabolism | -0.06 | 0.18 |
| 2-aminobutyrate | Methionine, Cysteine, SAM and Taurine Metabolism | -0.05 | 0.33 |
| phytanate | Food Component/Plant | -0.05 | 0.03 |
| 1-palmitoyl-2-oleoyl-GPE (16:0/18:1) | Phospholipid Metabolism | -0.05 | 0.41 |
| ursodeoxycholate | Secondary Bile Acid Metabolism | -0.05 | 0.21 |
| eicosapentaenoate (EPA; 20:5n3) | Polyunsaturated Fatty Acid (n3 and n6) | -0.05 | 0.19 |
| serotonin | Tryptophan Metabolism | -0.05 | -0.17 |
| theanine | Food Component/Plant | -0.05 | 0.54 |
| 1-stearoyl-2-linoleoyl-GPE (18:0/18:2)* | Phospholipid Metabolism | -0.05 | 0.43 |
| 10-undecenoate (11:1n1) | Medium Chain Fatty Acid | -0.05 | 0.08 |
| palmitate (16:0) | Long Chain Fatty Acid | -0.05 | 0.23 |
| 1-(1-enyl-stearoyl)-2-linoleoyl-GPE (P-18:0/18:2)* | Plasmalogen | -0.04 | 0.51 |
| 3-hydroxyoctanoate | Fatty Acid, Monohydroxy | -0.04 | 0.59 |
| margarate (17:0) | Long Chain Fatty Acid | -0.04 | 0.28 |
| hexanoylcarnitine (C6) | Fatty Acid Metabolism(Acyl Carnitine) | -0.04 | 0.51 |
| 1-oleoyl-GPC (18:1) | Lysolipid | -0.04 | -0.09 |
| 10-heptadecenoate (17:1n7) | Long Chain Fatty Acid | -0.04 | 0.25 |
| sphinganine | Sphingolipid Metabolism | -0.04 | -0.19 |
| linolenoylcarnitine (C18:3)* | Fatty Acid Metabolism(Acyl Carnitine) | -0.04 | -0.08 |
| vanillylmandelate (VMA) | Phenylalanine and Tyrosine Metabolism | -0.04 | 0.04 |
| isoleucine | Leucine, Isoleucine and Valine Metabolism | -0.04 | -0.17 |
| gamma-glutamylglycine | Gamma-glutamyl Amino Acid | -0.03 | -0.30 |
| palmitoyl-docosahexaenoyl-glycerol (16:0/22:6) [1]* | Diacylglycerol | -0.03 | 0.38 |
| beta-cryptoxanthin | Food Component/Plant | -0.03 | 0.15 |
| dodecanedioate | Fatty Acid, Dicarboxylate | -0.03 | 0.07 |
| sphingomyelin (d18:1/21:0, d17:1/22:0, d16:1/23:0)* | Sphingolipid Metabolism | -0.02 | 0.37 |
| 2,3-dihydroxyisovalerate | Food Component/Plant | -0.02 | -0.22 |
| 4-imidazoleacetate | Histidine Metabolism | -0.02 | -0.04 |
| oleoyl-linoleoyl-glycerol (18:1/18:2) [2] | Diacylglycerol | -0.02 | 0.30 |
| palmitoleoyl-arachidonoyl-glycerol (16:1/20:4) [2]* | Diacylglycerol | -0.02 | 0.13 |

| | | | |
|---|---|---|---|
| arginine | Urea cycle; Arginine and Proline Metabolism | -0.02 | 0.11 |
| 2-linoleoylglycerol (18:2) | Monoacylglycerol | -0.02 | -0.53 |
| sphingomyelin (d18:2/24:1, d18:1/24:2)* | Sphingolipid Metabolism | -0.02 | -0.14 |
| palmitoyl-linoleoyl-glycerol (16:0/18:2) [1]* | Diacylglycerol | -0.02 | 0.47 |
| pregnenolone sulfate | Steroid | -0.02 | -0.27 |
| 1-palmitoyl-2-alpha-linolenoyl-GPC (16:0/18:3n3)* | Phospholipid Metabolism | -0.02 | 0.19 |
| 3-hydroxybutyrylcarnitine (1) | Fatty Acid Metabolism(Acyl Carnitine) | -0.02 | 0.52 |
| 1-(1-enyl-stearoyl)-2-oleoyl-GPE (P-18:0/18:1) | Plasmalogen | -0.01 | 0.51 |
| phenylpyruvate | Phenylalanine and Tyrosine Metabolism | -0.01 | -0.24 |
| 3-methylhistidine | Histidine Metabolism | -0.01 | 0.20 |
| stearoyl-arachidonoyl-glycerol (18:0/20:4) [1]* | Diacylglycerol | -0.01 | 0.50 |
| caprate (10:0) | Medium Chain Fatty Acid | -0.01 | 0.49 |
| oleoyl-linoleoyl-glycerol (18:1/18:2) [1] | Diacylglycerol | -0.01 | 0.28 |
| N-acetylkynurenine (2) | Tryptophan Metabolism | -0.01 | 0.09 |
| 1-palmitoyl-2-arachidonoyl-GPC (16:0/20:4n6) | Phospholipid Metabolism | -0.01 | 0.02 |
| palmitoyl-arachidonoyl-glycerol (16:0/20:4) [1]* | Diacylglycerol | 0.00 | 0.48 |
| 21-hydroxypregnenolone disulfate | Steroid | 0.00 | -0.20 |
| umbelliferone sulfate | Food Component/Plant | 0.00 | 0.01 |

**Supplementary Table 5**. Pearson correlation coefficient between IL-1ra and all measured metabolites in the Null condition for both LTBI and TB.

| TNFa signature |
| --- |
| C3 |
| CCL4 |
| CD44 |
| CD83 |
| IRAK2 |
| NFKB1 |
| NFKB2 |
| NFKBIA |
| NFKBIZ |
| POU2F2 |
| RELB |
| SOCS3 |
| SRC |
| TNFAIP3 |

**Supplementary Table 6**. Unique cytokine-induced gene signature derived from TNF single cytokine stimulation as previously described[15].

| Gene | p-value | q-value | Fold change |
|---|---|---|---|
| IL1A | 3.43E-31 | 2.09E-28 | 212.311701 |
| CCL20 | 3.54E-29 | 7.20E-27 | 300.568543 |
| IL6 | 2.95E-29 | 7.20E-27 | 136.070461 |
| IL1B | 1.39E-28 | 2.13E-26 | 84.2357019 |
| CCL19 | 4.53E-25 | 5.52E-23 | 23.7958806 |
| IL10 | 2.52E-24 | 2.56E-22 | 22.4348548 |
| CCL2 | 4.46E-22 | 3.89E-20 | 69.4070409 |
| CXCL2 | 6.80E-21 | 5.19E-19 | 17.6122855 |
| CXCL1 | 5.58E-20 | 3.78E-18 | 7.24472043 |
| C3 | 2.52E-19 | 1.54E-17 | 25.3160454 |
| CCL3 | 3.31E-19 | 1.84E-17 | 12.3458144 |
| BATF3 | 7.24E-18 | 3.68E-16 | 10.3338066 |
| CCL4 | 8.44E-18 | 3.96E-16 | 5.72422376 |
| LILRB1 | 1.10E-16 | 4.80E-15 | 3.90816135 |
| SOCS3 | 9.14E-16 | 3.72E-14 | 5.15177685 |
| BATF | 2.88E-15 | 1.10E-13 | 3.41543669 |
| ADA | 5.79E-14 | 1.96E-12 | 2.7834998 |
| CD80 | 5.78E-14 | 1.96E-12 | 4.43455667 |
| TNFSF15 | 1.98E-13 | 6.36E-12 | 16.4780084 |
| SELPLG | 2.90E-13 | 8.86E-12 | 0.37229031 |
| IRAK2 | 5.89E-13 | 1.71E-11 | 6.46316131 |
| LAMP3 | 8.68E-13 | 2.41E-11 | 5.92884047 |
| PTGS2 | 1.04E-12 | 2.76E-11 | 14.6649475 |
| LAIR1 | 2.07E-12 | 5.25E-11 | 2.13989502 |
| POU2F2 | 3.79E-12 | 9.24E-11 | 2.26164102 |
| CD1D | 1.04E-11 | 2.45E-10 | 0.2482714 |
| TNFAIP6 | 1.12E-11 | 2.53E-10 | 4.95347156 |
| CCL22 | 1.36E-11 | 2.96E-10 | 7.0972002 |
| NFKBIZ | 1.47E-11 | 3.10E-10 | 3.13342401 |
| NFKB1 | 1.78E-11 | 3.61E-10 | 3.07008809 |
| EBI3 | 2.23E-11 | 4.39E-10 | 2.84525892 |
| LILRB4 | 2.58E-11 | 4.91E-10 | 4.50950805 |
| TNFRSF9 | 4.24E-11 | 7.84E-10 | 4.03643003 |
| SRC | 8.78E-11 | 1.57E-09 | 4.238783 |
| MSR1 | 9.04E-11 | 1.58E-09 | 0.24770417 |
| LILRA3 | 2.29E-10 | 3.87E-09 | 2.91298853 |
| SLAMF7 | 4.28E-10 | 7.05E-09 | 2.76580607 |
| CD40LG | 1.66E-09 | 2.66E-08 | 0.57192069 |
| TNF | 4.27E-09 | 6.69E-08 | 2.29017836 |
| FCER1A | 5.09E-09 | 7.76E-08 | 0.23207131 |
| CD14 | 5.23E-09 | 7.77E-08 | 2.14369557 |
| CLEC5A | 6.73E-09 | 9.55E-08 | 4.81805219 |

| | | | |
|---|---|---|---|
| TRAF1 | 6.62E-09 | 9.55E-08 | 2.03390181 |
| CCR2 | 7.78E-09 | 1.08E-07 | 0.33110713 |
| IL2RA | 7.98E-09 | 1.08E-07 | 4.1747798 |
| CCL7 | 1.22E-08 | 1.62E-07 | 5.64608199 |
| DUSP4 | 1.39E-08 | 1.81E-07 | 4.29974158 |
| IL23A | 2.00E-08 | 2.54E-07 | 4.27694314 |
| NFKBIA | 2.07E-08 | 2.58E-07 | 2.57754936 |
| IL8 | 2.14E-08 | 2.62E-07 | 3.44525325 |
| AHR | 2.74E-08 | 3.27E-07 | 1.55085627 |
| ALDH1A1 | 5.11E-08 | 5.99E-07 | 0.31581903 |
| ICAM5 | 9.11E-08 | 1.05E-06 | 3.73567785 |
| CXCL13 | 1.22E-07 | 1.35E-06 | 2.24874481 |
| IRAK3 | 1.22E-07 | 1.35E-06 | 2.88747799 |
| CD274 | 1.64E-07 | 1.79E-06 | 6.21347375 |
| CDKN1A | 1.67E-07 | 1.79E-06 | 3.60580151 |
| PDCD1LG2 | 4.50E-07 | 4.74E-06 | 3.24891949 |
| KCNJ2 | 7.52E-07 | 7.78E-06 | 3.48865275 |
| CASP8 | 9.28E-07 | 9.44E-06 | 0.60129254 |
| PLAU | 1.04E-06 | 1.04E-05 | 7.67171924 |
| RELB | 1.72E-06 | 1.70E-05 | 2.1874802 |
| CCRL2 | 1.86E-06 | 1.78E-05 | 3.64517996 |
| CD40 | 1.87E-06 | 1.78E-05 | 2.21657161 |
| TNFAIP3 | 2.13E-06 | 2.00E-05 | 2.01128837 |
| TNFRSF4 | 2.23E-06 | 2.06E-05 | 2.53379536 |
| TNFSF12 | 3.45E-06 | 3.14E-05 | 0.82198616 |
| CASP2 | 3.94E-06 | 3.53E-05 | 0.77089325 |
| CD46 | 4.33E-06 | 3.82E-05 | 0.60245566 |
| IL7R | 5.78E-06 | 4.96E-05 | 0.72868945 |
| NFATC3 | 5.78E-06 | 4.96E-05 | 0.72909413 |
| HLA.DMB | 5.86E-06 | 4.97E-05 | 0.72039147 |
| ICAM1 | 6.12E-06 | 5.11E-05 | 2.35325886 |
| JAK3 | 6.48E-06 | 5.34E-05 | 1.63910012 |
| IL16 | 6.67E-06 | 5.42E-05 | 0.68323606 |
| MARCO | 7.47E-06 | 6.00E-05 | 5.50605871 |
| CSF3R | 7.69E-06 | 6.09E-05 | 0.51794721 |
| SELL | 8.03E-06 | 6.28E-05 | 0.6690134 |
| PYCARD | 9.99E-06 | 7.72E-05 | 0.5088453 |
| BID | 1.18E-05 | 8.99E-05 | 2.97096169 |
| S1PR1 | 1.59E-05 | 0.00011975 | 0.69552438 |
| ITGA4 | 1.81E-05 | 0.00013479 | 0.76409875 |
| IDO1 | 1.85E-05 | 0.00013584 | 2.51373061 |
| CXCR2 | 2.25E-05 | 0.00016354 | 0.52178957 |
| LILRA1 | 2.45E-05 | 0.00017617 | 2.45722501 |

| | | | |
|---|---|---|---|
| IKBKE | 2.63E-05 | 0.0001865 | 1.53738768 |
| CYB561 | 3.04E-05 | 0.00021345 | 0.66560952 |
| KLF2 | 3.60E-05 | 0.00024923 | 0.66820886 |
| TICAM1 | 4.13E-05 | 0.00028295 | 1.97800218 |
| CCL23 | 4.35E-05 | 0.00029163 | 2.11730627 |
| CYBB | 4.35E-05 | 0.00029163 | 1.4926294 |
| IL1RN | 4.83E-05 | 0.00031993 | 3.74745204 |
| NFKB2 | 4.89E-05 | 0.00032091 | 1.91395377 |
| CD36 | 5.50E-05 | 0.00035679 | 0.58312416 |
| ATG7 | 5.86E-05 | 0.00037657 | 1.87104598 |
| SELE | 6.10E-05 | 0.00038737 | 1.53308961 |
| FER1L3 | 6.28E-05 | 0.00039472 | 2.6818528 |
| CTSS | 7.77E-05 | 0.0004836 | 0.64444716 |
| CD83 | 8.07E-05 | 0.0004923 | 1.86846557 |
| IFIT2 | 8.07E-05 | 0.0004923 | 0.45531283 |
| CISH | 9.29E-05 | 0.00056106 | 2.81329332 |
| GAS6 | 9.50E-05 | 0.00056824 | 1.47604225 |
| CD244 | 0.00010707 | 0.0006341 | 0.72778338 |
| BCL3 | 0.00012504 | 0.00073339 | 2.07021415 |
| HLA.DMA | 0.00012737 | 0.00073998 | 0.72211274 |
| CCND3 | 0.00014055 | 0.00080884 | 0.77631712 |
| SIGIRR | 0.00017294 | 0.00098592 | 0.78599771 |
| CASP10 | 0.00019282 | 0.00108908 | 1.85097323 |
| LTB4R2 | 0.00019467 | 0.00108943 | 0.63438784 |
| CLEC7A | 0.00022668 | 0.00125702 | 0.6194544 |
| TNFSF10 | 0.00029406 | 0.00161602 | 0.61687154 |
| CCL18 | 0.00029984 | 0.00163304 | 1.43183676 |
| PTAFR | 0.00031731 | 0.00171291 | 1.62913479 |
| CD82 | 0.00036271 | 0.00194083 | 1.80731 |
| IL23R | 0.00036593 | 0.00194101 | 2.08019679 |
| CFD | 0.00039919 | 0.00209919 | 0.41915334 |
| ITGAL | 0.00051134 | 0.00266596 | 0.71677539 |
| CD96 | 0.00061903 | 0.00320007 | 0.78448023 |
| ICOSLG | 0.00065448 | 0.00335488 | 1.8685809 |
| SPP1 | 0.00066704 | 0.00339079 | 5.24273659 |
| CARD9 | 0.00071088 | 0.00358379 | 1.89067577 |
| CXCL10 | 0.00071791 | 0.00358957 | 2.13308281 |
| HPSE | 0.00073118 | 0.00362619 | 2.09786235 |
| HAVCR2 | 0.00075553 | 0.00367935 | 1.96191698 |
| TGFBR2 | 0.00075583 | 0.00367935 | 0.72573864 |
| VAMP5 | 0.00076 | 0.00367935 | 0.75889806 |
| LCK | 0.00079416 | 0.00381445 | 0.87109985 |
| CD27 | 0.00085273 | 0.00406377 | 0.75457751 |

| | | | |
|---|---|---|---|
| FCGRT | 0.00089073 | 0.00421198 | 0.73925756 |
| SOCS1 | 0.0009399 | 0.00441031 | 2.00084581 |
| CD3E | 0.00129682 | 0.00603863 | 0.77590935 |
| LAG3 | 0.00140841 | 0.00650856 | 1.90832121 |
| PECAM1 | 0.00150262 | 0.00689172 | 0.71562813 |
| LHFPL2 | 0.00152647 | 0.00694886 | 2.37025136 |
| KLRK1 | 0.0015997 | 0.00722827 | 0.81133409 |
| CD22 | 0.00177202 | 0.00794803 | 2.39445935 |
| ICAM3 | 0.0019465 | 0.0086669 | 0.74639282 |
| IL1R2 | 0.00200718 | 0.00887232 | 1.76261942 |
| CSF2 | 0.00229442 | 0.01006904 | 1.22930763 |
| CCL5 | 0.0026367 | 0.01148848 | 0.74536035 |
| CLEC6A | 0.00274293 | 0.0118462 | 2.10609416 |
| KLRB1 | 0.00275764 | 0.0118462 | 0.78962194 |
| IGF2R | 0.00297059 | 0.01267175 | 0.75002209 |
| CD4 | 0.00300011 | 0.0127088 | 0.82125491 |
| ABCB1 | 0.00307494 | 0.01293595 | 0.8313858 |
| POLR1B | 0.00319791 | 0.01336113 | 1.8567145 |
| IL1R1 | 0.00324724 | 0.01341943 | 2.66620929 |
| TLR2 | 0.00327786 | 0.01341943 | 1.55679825 |
| sCTLA4 | 0.00327617 | 0.01341943 | 1.97778007 |
| CD163 | 0.00335888 | 0.01365945 | 1.57298181 |
| TLR9 | 0.00363583 | 0.01468779 | 0.56050961 |
| IRF8 | 0.00368463 | 0.014787 | 0.7829239 |
| FCAR | 0.00385377 | 0.0153647 | 1.38541194 |
| C1QB | 0.00416485 | 0.01628563 | 1.7231377 |
| IFNA1.13 | 0.00415787 | 0.01628563 | 1.22245725 |
| MBP | 0.0041368 | 0.01628563 | 0.79074532 |
| CD247 | 0.00429836 | 0.01659493 | 0.85589517 |
| ICAM2 | 0.00427783 | 0.01659493 | 0.89108688 |
| STAT6 | 0.00442326 | 0.01696974 | 0.70405303 |
| ETS1 | 0.00461676 | 0.0176014 | 0.82120368 |
| TRAF3 | 0.00468297 | 0.01774293 | 1.44733713 |
| ZAP70 | 0.00475 | 0.0178858 | 0.81264434 |
| GZMA | 0.00484075 | 0.01800523 | 0.75953587 |
| PSMB10 | 0.00481794 | 0.01800523 | 0.84883497 |
| TCF7 | 0.00524422 | 0.01938772 | 0.80390393 |
| IFNG | 0.00571594 | 0.02075431 | 1.21169367 |
| NLRP3 | 0.00565143 | 0.02075431 | 1.64264762 |
| TNFSF8 | 0.00570429 | 0.02075431 | 1.46439223 |
| CD5 | 0.00575757 | 0.02075554 | 0.82957309 |
| ITGA6 | 0.00578433 | 0.02075554 | 1.18908924 |
| FYN | 0.00603091 | 0.02151377 | 0.7897243 |

| | | | |
|---|---|---|---|
| MAP4K1 | 0.00640358 | 0.02244933 | 0.85128965 |
| MAPK11 | 0.00638657 | 0.02244933 | 0.57492036 |
| SH2D1A | 0.00634492 | 0.02244933 | 0.82758476 |
| TBX21 | 0.00649259 | 0.02263131 | 0.81782609 |
| FCGR3A.B | 0.00655248 | 0.0227103 | 0.65242591 |
| PLAUR | 0.00673515 | 0.02321153 | 1.90470689 |
| IFNAR2 | 0.00679705 | 0.02329326 | 0.73547948 |
| TRAF2 | 0.00737003 | 0.02511574 | 1.71960001 |
| CXCR3 | 0.0075181 | 0.02537897 | 0.81245398 |
| IL17F | 0.00753048 | 0.02537897 | 1.10094317 |
| NCF4 | 0.00771893 | 0.02587114 | 0.73172274 |
| ENTPD1 | 0.00797935 | 0.02659783 | 0.66945316 |
| NOTCH1 | 0.0080744 | 0.02676839 | 0.76211732 |
| LEF1 | 0.00879035 | 0.0289844 | 0.89700691 |
| CSF2RB | 0.00902795 | 0.02950632 | 1.40375874 |
| CXCL9 | 0.00909375 | 0.02950632 | 1.33990527 |
| MAP4K2 | 0.00906815 | 0.02950632 | 0.82301292 |
| CCR7 | 0.00931243 | 0.03005599 | 1.25983431 |
| CD9 | 0.0103004 | 0.03306971 | 0.58875487 |
| GBP2 | 0.0105318 | 0.03363559 | 0.69579201 |
| IL12RB1 | 0.011472 | 0.0364475 | 0.60897927 |
| SMAD3 | 0.0117892 | 0.0372612 | 1.28071991 |
| ARHGDIB | 0.0120456 | 0.03787534 | 0.85433097 |
| TRAFD1 | 0.0121496 | 0.03800644 | 0.74259937 |
| PML | 0.0123071 | 0.03830271 | 0.76551789 |
| LTBR | 0.012402 | 0.03840213 | 2.00848815 |
| IL6R | 0.0125199 | 0.03857141 | 0.75910587 |
| CD209 | 0.0132276 | 0.04053603 | 1.37714933 |
| FCER1G | 0.0132905 | 0.04053603 | 1.49507825 |
| IL12B | 0.0136527 | 0.04143357 | 1.22335744 |
| MAPK1 | 0.0137494 | 0.04152047 | 0.80608116 |
| HLA.C | 0.0138663 | 0.04166721 | 0.71163926 |
| KREMEN1 | 0.0142776 | 0.04269282 | 2.14163102 |
| NFATC2 | 0.014464 | 0.04303922 | 0.80822451 |
| PPBP | 0.0147094 | 0.04355696 | 0.75049637 |
| CXCR1 | 0.0154676 | 0.04558085 | 0.67646679 |
| SDHA | 0.015728 | 0.04612538 | 0.87478234 |
| ARG2 | 0.0158037 | 0.04612563 | 1.61150088 |
| ABL1 | 0.0171857 | 0.04992037 | 0.82545473 |
| CD99 | 0.0179527 | 0.05190117 | 0.8267941 |
| MR1 | 0.0181272 | 0.05215845 | 1.67273741 |
| CCR1 | 0.0182493 | 0.05226325 | 1.67253452 |
| CD28 | 0.0189324 | 0.05371518 | 0.86730721 |

| | | | |
|---|---|---|---|
| NOD1 | 0.0189039 | 0.05371518 | 0.71072877 |
| TRAF4 | 0.0194128 | 0.05482319 | 1.80643831 |
| IL21R | 0.0195709 | 0.05501497 | 1.58548255 |
| BCL2L11 | 0.0206973 | 0.05791446 | 0.88999183 |
| CTLA4.TM | 0.0214969 | 0.05987721 | 1.80442979 |
| CD44 | 0.0219813 | 0.06091361 | 1.29736012 |
| FAS | 0.0220687 | 0.06091361 | 0.71111214 |
| BCL2 | 0.0225096 | 0.0618507 | 0.91879697 |
| CLEC4A | 0.0228753 | 0.06257369 | 1.44631318 |
| DEFB1 | 0.0236658 | 0.06416061 | 0.82098571 |
| TRAF5 | 0.0236122 | 0.06416061 | 0.9492829 |
| PAX5 | 0.0245032 | 0.0658456 | 0.60878469 |
| Sep-04 | 0.024423 | 0.0658456 | 0.57727473 |
| CD79B | 0.025349 | 0.06781969 | 0.7502774 |
| DEFA1 | 0.0261553 | 0.06967132 | 0.68171377 |
| ICOS | 0.02676 | 0.07066494 | 1.54661702 |
| KLRC2 | 0.026656 | 0.07066494 | 0.86731502 |
| IL32 | 0.0280509 | 0.07343798 | 0.67748363 |
| JAK2 | 0.0280509 | 0.07343798 | 0.79497257 |
| CCR10 | 0.0285155 | 0.07433528 | 0.89192358 |
| JAK1 | 0.0289307 | 0.0747785 | 0.82202888 |
| RARRES3 | 0.0289307 | 0.0747785 | 0.94118589 |
| CD3D | 0.0298342 | 0.07675927 | 0.84863318 |
| TNFRSF10C | 0.0299487 | 0.07675927 | 0.75268911 |
| BTLA | 0.030411 | 0.07761803 | 0.87786804 |
| HLA.DPB1 | 0.031956 | 0.0812215 | 0.81986655 |
| TNFRSF8 | 0.0342048 | 0.08657646 | 1.70489041 |
| LILRA2 | 0.0349853 | 0.08818609 | 0.82799503 |
| CRADD | 0.0358408 | 0.08997073 | 1.62697492 |
| PDCD2 | 0.0361369 | 0.08997351 | 0.79849596 |
| SMARCD3 | 0.0359919 | 0.08997351 | 0.58801709 |
| MS4A1 | 0.0370005 | 0.09174921 | 0.83399636 |
| XCR1 | 0.0371854 | 0.09183439 | 0.81753477 |
| CEBPB | 0.0376939 | 0.09271483 | 1.36084524 |
| CD74 | 0.0388263 | 0.09511664 | 0.82851689 |
| LTB4R | 0.0401344 | 0.09792794 | 0.79819602 |
| CREB5 | 0.0419357 | 0.10191545 | 0.70124622 |
| CD276 | 0.0431552 | 0.10405009 | 1.59065727 |
| IL28A.B | 0.0430355 | 0.10405009 | 0.94951237 |
| SLC2A1 | 0.0448919 | 0.10781126 | 1.70450996 |
| IL13RA1 | 0.0455834 | 0.10904264 | 0.82272604 |
| SMAD5 | 0.0464061 | 0.11057704 | 1.05105911 |
| CLEC4E | 0.0472501 | 0.11215004 | 1.35900616 |

| | | | |
|---|---|---|---|
| NT5E | 0.048583 | 0.11486678 | 1.67988098 |
| TLR1 | 0.048968 | 0.11533004 | 0.80880735 |
| IRF1 | 0.0500236 | 0.11736306 | 0.78108095 |
| CX3CR1 | 0.0509164 | 0.11900002 | 0.79734942 |
| CCL8 | 0.0512057 | 0.11921938 | 1.76455938 |
| KIT | 0.0518135 | 0.1201758 | 0.80363649 |
| SLAMF6 | 0.0528695 | 0.12216059 | 0.81562393 |
| BTK | 0.0540587 | 0.12435033 | 0.84838205 |
| KLRG2 | 0.0542249 | 0.12435033 | 0.86862598 |
| IRF3 | 0.0561793 | 0.12787079 | 0.73040776 |
| KLRG1 | 0.056164 | 0.12787079 | 0.70173295 |
| CD58 | 0.057559 | 0.13052413 | 1.23379696 |
| PTK2 | 0.0583616 | 0.13185399 | 0.83897981 |
| IFNAR1 | 0.0597879 | 0.13457793 | 1.34815558 |
| APOL6 | 0.0612457 | 0.13735249 | 0.79885911 |
| HLA.A | 0.0616672 | 0.13779118 | 0.83238104 |
| CD55 | 0.0624808 | 0.13909959 | 0.69885206 |
| CSF1R | 0.0630671 | 0.13938743 | 1.65715666 |
| STAT5B | 0.062946 | 0.13938743 | 0.82923964 |
| MME | 0.0633755 | 0.13956338 | 0.66101871 |
| GBP5 | 0.0655693 | 0.14387508 | 0.70290566 |
| CD24 | 0.0669112 | 0.1457778 | 0.88280579 |
| PSMB8 | 0.0669144 | 0.1457778 | 0.84504379 |
| PSMB5 | 0.0682731 | 0.14820851 | 1.58611787 |
| PSMB9 | 0.0701414 | 0.1517243 | 0.81242357 |
| TNFRSF13C | 0.0730081 | 0.15736728 | 0.88096707 |
| CAMP | 0.073963 | 0.15886419 | 0.688927 |
| STAT2 | 0.0769787 | 0.16476143 | 0.86756094 |
| CASP1 | 0.0805956 | 0.17070638 | 1.23722937 |
| TLR8 | 0.0800706 | 0.17070638 | 0.75788192 |
| TMEM173 | 0.0805958 | 0.17070638 | 0.94737703 |
| IKBKAP | 0.0810036 | 0.17097646 | 0.94802598 |
| ATM | 0.0819438 | 0.17236454 | 0.71593722 |
| GBP4 | 0.0838049 | 0.1756735 | 0.80465764 |
| CEACAM1 | 0.0841446 | 0.17578153 | 1.64022532 |
| KLRF2 | 0.0844598 | 0.17583781 | 1.07387644 |
| C4A.B | 0.0850947 | 0.17655703 | 0.66902777 |
| IL10RA | 0.0879606 | 0.18188463 | 0.86882228 |
| APOL1 | 0.0916957 | 0.18644833 | 0.87767821 |
| CD79A | 0.0911131 | 0.18644833 | 0.85576525 |
| CD81 | 0.0916959 | 0.18644833 | 1.23170093 |
| KLRC4 | 0.0916545 | 0.18644833 | 0.82044073 |
| LILRB2 | 0.0911131 | 0.18644833 | 1.30401729 |

| | | | |
|---|---|---|---|
| NCAM1 | 0.0922502 | 0.18695223 | 0.75256442 |
| ETV7 | 0.0992554 | 0.20048276 | 0.63763115 |
| CD45RB | 0.103348 | 0.20806033 | 1.15826792 |
| BCL6 | 0.106602 | 0.21390533 | 1.33763732 |
| RAF1 | 0.109264 | 0.21781386 | 0.82334329 |
| TIGIT | 0.109264 | 0.21781386 | 0.89377029 |
| ITGB2 | 0.111978 | 0.222497 | 0.83965874 |
| TGFB1 | 0.113355 | 0.22450179 | 0.82002967 |
| IL6ST | 0.114396 | 0.22583029 | 1.15947925 |
| CTNNB1 | 0.115445 | 0.22716597 | 1.15382306 |
| IL1RL1 | 0.118917 | 0.23259965 | 1.51005487 |
| TNFSF13B | 0.118969 | 0.23259965 | 0.98631139 |
| MALT1 | 0.124109 | 0.24187377 | 0.91407852 |
| PRF1 | 0.126352 | 0.24546089 | 0.84738765 |
| CD53 | 0.128627 | 0.24908721 | 1.24482765 |
| CMKLR1 | 0.131754 | 0.25419681 | 0.69474287 |
| HLA.DRA | 0.132099 | 0.25419681 | 1.13061755 |
| C14orf166 | 0.135246 | 0.25943415 | 0.90161911 |
| PSMC2 | 0.137242 | 0.26243768 | 1.21821942 |
| IL7 | 0.139108 | 0.26517462 | 0.88131948 |
| HLA.B | 0.139667 | 0.26541081 | 0.84691847 |
| AIRE | 0.14086 | 0.26684658 | 0.93640828 |
| CD6 | 0.142125 | 0.26758102 | 0.90415812 |
| MIF | 0.142125 | 0.26758102 | 1.34301833 |
| CCR5 | 0.143762 | 0.26941043 | 1.23552826 |
| LIF | 0.14398 | 0.26941043 | 1.81056879 |
| IFITM1 | 0.148414 | 0.27601384 | 1.18226054 |
| IRAK1 | 0.147986 | 0.27601384 | 1.43674796 |
| DUSP3 | 0.150558 | 0.27915009 | 1.30344798 |
| DEFB4A | 0.156091 | 0.28593246 | 0.96618583 |
| IL20 | 0.156091 | 0.28593246 | 0.97400494 |
| RAG1 | 0.156091 | 0.28593246 | 0.97013566 |
| THY1 | 0.156091 | 0.28593246 | 0.97783239 |
| AICDA | 0.160932 | 0.29079776 | 1.05879288 |
| CXCL11 | 0.160932 | 0.29079776 | 1.06059173 |
| FOXP3 | 0.162055 | 0.29079776 | 1.07763529 |
| IRGM | 0.159769 | 0.29079776 | 1.02710324 |
| ITGAE | 0.162031 | 0.29079776 | 0.98535241 |
| NCR1 | 0.162084 | 0.29079776 | 0.89954789 |
| SKI | 0.160269 | 0.29079776 | 0.93711201 |
| CD8B | 0.164511 | 0.29428654 | 0.92099741 |
| CD19 | 0.169493 | 0.29969919 | 0.74935642 |
| CR1 | 0.168561 | 0.29969919 | 0.87422952 |

| | | | |
|---|---|---|---|
| HLA.DPA1 | 0.169502 | 0.29969919 | 0.87315519 |
| IKBKG | 0.168561 | 0.29969919 | 0.94193082 |
| VTN | 0.170707 | 0.30095743 | 0.92160854 |
| SCARF1 | 0.171381 | 0.30127496 | 1.45139378 |
| TOLLIP | 0.174265 | 0.30546451 | 0.94469799 |
| ITLN1 | 0.176005 | 0.30763052 | 1.1476884 |
| GNLY | 0.179619 | 0.31305026 | 0.88972414 |
| EGR2 | 0.180875 | 0.31434117 | 1.44450382 |
| PDCD1 | 0.186736 | 0.323605 | 0.84438391 |
| EDNRB | 0.188065 | 0.32498484 | 1.34089848 |
| S100A8 | 0.190179 | 0.32770958 | 1.18120961 |
| CSF1 | 0.193623 | 0.33270431 | 0.87894208 |
| PPIA | 0.199035 | 0.34103272 | 1.25074439 |
| STAT1 | 0.199588 | 0.34103272 | 0.82554514 |
| IL4R | 0.203877 | 0.34738818 | 1.23528163 |
| FKBP5 | 0.20496 | 0.34821003 | 0.88890059 |
| GFI1 | 0.205501 | 0.34821003 | 0.94422952 |
| TYK2 | 0.206592 | 0.34908898 | 0.9034715 |
| KLRC3 | 0.209884 | 0.35367193 | 0.8366476 |
| LILRB3 | 0.216018 | 0.36300545 | 0.89537441 |
| FCGR2B | 0.217147 | 0.36390019 | 0.83392641 |
| CD2 | 0.219988 | 0.36765118 | 0.94258158 |
| TAP2 | 0.221706 | 0.36945722 | 0.88409787 |
| TNFRSF1B | 0.22228 | 0.36945722 | 1.19749032 |
| TP53 | 0.223432 | 0.37036283 | 1.09566746 |
| BCAP31 | 0.232212 | 0.38272745 | 0.95361954 |
| C8G | 0.23338 | 0.38272745 | 0.93350594 |
| IL4 | 0.232643 | 0.38272745 | 0.91201408 |
| PRKCD | 0.233401 | 0.38272745 | 1.25379718 |
| LTF | 0.23699 | 0.38757078 | 0.79073599 |
| CD3EAP | 0.24763 | 0.40291069 | 0.95418499 |
| LILRA6 | 0.247691 | 0.40291069 | 1.91359298 |
| CXCR4 | 0.250498 | 0.40639303 | 0.86551876 |
| B2M | 0.254273 | 0.41033474 | 0.90280042 |
| TNFRSF11A | 0.254201 | 0.41033474 | 0.74298192 |
| STAT4 | 0.264531 | 0.4257623 | 0.93657972 |
| IKZF2 | 0.271007 | 0.43389572 | 1.12282259 |
| RORC | 0.27048 | 0.43389572 | 0.8406633 |
| ATG5 | 0.278415 | 0.4445894 | 1.34550337 |
| DPP4 | 0.289657 | 0.4613336 | 0.97334206 |
| ATG16L1 | 0.292796 | 0.46511865 | 1.06504958 |
| CXCR6 | 0.295033 | 0.46647775 | 1.43213056 |
| IL13 | 0.295181 | 0.46647775 | 0.85225201 |

| | | | |
|---|---|---|---|
| GZMK | 0.296995 | 0.46813165 | 0.93233091 |
| CD164 | 0.301243 | 0.4723862 | 0.91049185 |
| NOD2 | 0.301243 | 0.4723862 | 1.29559876 |
| IL12A | 0.307525 | 0.48100064 | 0.97276741 |
| FN1 | 0.310438 | 0.48431504 | 0.82616852 |
| ANKRD22 | 0.31536 | 0.48847726 | 1.38419865 |
| BAX | 0.317864 | 0.48847726 | 0.94644869 |
| CCBP2 | 0.318713 | 0.48847726 | 1.0254983 |
| CD160 | 0.322005 | 0.48847726 | 0.86480094 |
| CD45RA | 0.324529 | 0.48847726 | 0.98905825 |
| CD97 | 0.326022 | 0.48847726 | 0.93570684 |
| CEACAM8 | 0.320801 | 0.48847726 | 0.74039445 |
| IL19 | 0.325767 | 0.48847726 | 1.00189353 |
| IL2 | 0.325767 | 0.48847726 | 1.034742 |
| IL22 | 0.325767 | 0.48847726 | 1.0306703 |
| IL26 | 0.320108 | 0.48847726 | 0.98735117 |
| MAPK14 | 0.327519 | 0.48847726 | 0.96051297 |
| MASP2 | 0.325767 | 0.48847726 | 1.00370646 |
| PIGR | 0.320108 | 0.48847726 | 0.99101555 |
| PLA2G2A | 0.320108 | 0.48847726 | 0.98075405 |
| RAG2 | 0.320108 | 0.48847726 | 0.99101555 |
| UBE2L3 | 0.32752 | 0.48847726 | 1.00654362 |
| VCAM1 | 0.324406 | 0.48847726 | 1.02351468 |
| TLR3 | 0.329242 | 0.48865601 | 1.0528462 |
| XBP1 | 0.329022 | 0.48865601 | 1.19473529 |
| TIRAP | 0.330712 | 0.48964641 | 0.84712689 |
| LILRA5 | 0.33204 | 0.49042228 | 1.14940163 |
| CASP4 | 0.335075 | 0.49225094 | 1.1406973 |
| IL1RAP | 0.335075 | 0.49225094 | 1.19323815 |
| TLR5 | 0.335699 | 0.49225094 | 0.83518517 |
| HLA.DQA1 | 0.34947 | 0.51121511 | 1.12992421 |
| LY96 | 0.360797 | 0.52517369 | 0.90837065 |
| NFATC1 | 0.361595 | 0.52517369 | 0.97400852 |
| TFRC | 0.361595 | 0.52517369 | 0.94366474 |
| RELA | 0.3648 | 0.52857007 | 1.20784221 |
| GUSB | 0.372888 | 0.53784839 | 0.96927289 |
| XCL1 | 0.372967 | 0.53784839 | 1.01761723 |
| C2 | 0.375812 | 0.54067292 | 1.2932077 |
| CIITA | 0.381735 | 0.547902 | 0.88459194 |
| ITGA2B | 0.385545 | 0.55207148 | 0.8647416 |
| CFP | 0.387724 | 0.55259729 | 1.11296868 |
| TAP1 | 0.387724 | 0.55259729 | 0.87414408 |
| GPI | 0.389394 | 0.55368378 | 1.16969954 |

| | | | |
|---|---|---|---|
| IL18RAP | 0.391068 | 0.55477088 | 1.16180907 |
| CFB | 0.396128 | 0.56007461 | 1.02194057 |
| IL28A | 0.396643 | 0.56007461 | 0.9261495 |
| PDGFB | 0.398913 | 0.56197905 | 1.07397991 |
| ATG12 | 0.41275 | 0.57893067 | 0.85002784 |
| CCL13 | 0.412844 | 0.57893067 | 1.02274331 |
| GATA3 | 0.417817 | 0.58411339 | 0.88218125 |
| TLR4 | 0.418455 | 0.58411339 | 0.97002504 |
| CEACAM6 | 0.420253 | 0.58528386 | 0.81710138 |
| IL2RG | 0.423713 | 0.58875838 | 1.15033494 |
| CCR6 | 0.42698 | 0.59194955 | 0.98689219 |
| KLRC1 | 0.428002 | 0.59202091 | 0.99556623 |
| TAGAP | 0.436131 | 0.60190025 | 0.98777742 |
| MCL1 | 0.437922 | 0.60300772 | 1.12754186 |
| CD70 | 0.442715 | 0.60686775 | 0.89480486 |
| IL11RA | 0.442156 | 0.60686775 | 0.81014778 |
| CHUK | 0.446033 | 0.6080529 | 0.91815778 |
| CTSG | 0.445258 | 0.6080529 | 0.88108677 |
| HRAS | 0.44657 | 0.6080529 | 1.20391874 |
| SLAMF1 | 0.468797 | 0.6368957 | 1.23558906 |
| TBP | 0.472742 | 0.63940714 | 1.01935516 |
| TGFBR1 | 0.471807 | 0.63940714 | 0.92797896 |
| HLA.DRB1 | 0.474617 | 0.64052294 | 1.12235184 |
| TNFRSF13B | 0.479907 | 0.64623238 | 1.18562766 |
| KIR3DL1 | 0.482274 | 0.6479893 | 1.08763539 |
| KLRF1 | 0.483841 | 0.64866596 | 1.05579116 |
| ITGAM | 0.484996 | 0.64878851 | 1.15067627 |
| IFIH1 | 0.487849 | 0.651177 | 1.19095673 |
| IL15 | 0.494111 | 0.65809544 | 0.99043431 |
| IRF5 | 0.500886 | 0.66473557 | 1.24508135 |
| RUNX1 | 0.501276 | 0.66473557 | 1.24443165 |
| IKZF1 | 0.504179 | 0.6671349 | 1.08122741 |
| LACTB | 0.507091 | 0.66953574 | 1.12421347 |
| HLA.DOB | 0.523646 | 0.68990078 | 1.31224854 |
| CALML4 | 0.531688 | 0.69898638 | 1.02623384 |
| STAT3 | 0.536685 | 0.70403839 | 1.06416569 |
| C4BPA | 0.548442 | 0.71791764 | 1.13857395 |
| BLNK | 0.551223 | 0.7189473 | 1.14149219 |
| C6 | 0.553943 | 0.7189473 | 1.173026 |
| NFIL3 | 0.552816 | 0.7189473 | 1.10844562 |
| ZBTB16 | 0.553341 | 0.7189473 | 0.98897092 |
| CCL15 | 0.563694 | 0.72542899 | 1.02126035 |
| CX3CL1 | 0.563694 | 0.72542899 | 1.02102012 |

| | | | |
|---|---|---|---|
| GP1BB | 0.562819 | 0.72542899 | 1.00555879 |
| IFI35 | 0.563215 | 0.72542899 | 1.28270996 |
| DEFB103A | 0.568237 | 0.72667625 | 0.99825116 |
| EOMES | 0.567464 | 0.72667625 | 0.89731847 |
| LGALS3 | 0.567123 | 0.72667625 | 1.20090831 |
| HLA.DRB3 | 0.579526 | 0.7380185 | 0.91746943 |
| SERPING1 | 0.578631 | 0.7380185 | 0.84752451 |
| BST2 | 0.584731 | 0.74155075 | 0.9955756 |
| TGFBI | 0.584731 | 0.74155075 | 0.90999782 |
| SYK | 0.587865 | 0.74397853 | 0.99163273 |
| TCF4 | 0.598361 | 0.75569402 | 0.95334857 |
| MAF | 0.603403 | 0.76048725 | 1.10522799 |
| PTPN2 | 0.607888 | 0.76142029 | 1.14269312 |
| PTPN22 | 0.607881 | 0.76142029 | 1.11340386 |
| TRAF6 | 0.607888 | 0.76142029 | 0.97447002 |
| LITAF | 0.612142 | 0.7651775 | 0.9596249 |
| ATG10 | 0.622806 | 0.77375084 | 1.18880163 |
| CD7 | 0.621761 | 0.77375084 | 1.19495062 |
| CR2 | 0.622305 | 0.77375084 | 1.07204763 |
| LTA | 0.628935 | 0.77977713 | 1.13401051 |
| LILRA4 | 0.635623 | 0.78647065 | 0.89674518 |
| S100A9 | 0.643367 | 0.79444103 | 1.08515866 |
| HLA.DQB1 | 0.646359 | 0.79652321 | 1.12634751 |
| ILF3 | 0.647726 | 0.79659851 | 1.02292048 |
| CFH | 0.657738 | 0.80404846 | 0.85253738 |
| GPR183 | 0.657576 | 0.80404846 | 1.12008716 |
| IRF7 | 0.656479 | 0.80404846 | 1.01799862 |
| C5 | 0.661618 | 0.80717396 | 0.91560548 |
| C1QA | 0.664184 | 0.80868711 | 1.02361437 |
| ICAM4 | 0.675353 | 0.82064807 | 0.92435504 |
| CD86 | 0.678564 | 0.82291062 | 1.18018743 |
| FADD | 0.68175 | 0.82503285 | 0.90721852 |
| MAP4K4 | 0.683019 | 0.82503285 | 1.1516657 |
| CASP3 | 0.68748 | 0.82714556 | 1.00903177 |
| CTSC | 0.686365 | 0.82714556 | 1.05312844 |
| BST1 | 0.691955 | 0.83089085 | 1.02081257 |
| IFI16 | 0.698686 | 0.83568325 | 0.98304134 |
| PTPN6 | 0.698686 | 0.83568325 | 0.9633485 |
| NOTCH2 | 0.700935 | 0.83673258 | 1.01019156 |
| C1QBP | 0.703186 | 0.8377802 | 1.0741087 |
| FCGR1A.B | 0.708183 | 0.84208895 | 1.17109816 |
| ACTA2 | 0.715336 | 0.8479308 | 1.01648195 |
| PPARG | 0.715876 | 0.8479308 | 0.92670496 |

| | | | |
|---|---|---|---|
| CD59 | 0.725842 | 0.85806903 | 1.04673214 |
| KCNJ15 | 0.733824 | 0.86582716 | 1.23034166 |
| ARG1 | 0.764021 | 0.89971585 | 0.86905718 |
| FCGR2A | 0.771861 | 0.90371441 | 0.97498383 |
| IKBKB | 0.76948 | 0.90371441 | 1.07116186 |
| PTPRC_all | 0.771861 | 0.90371441 | 0.96544241 |
| IL1RL2 | 0.776761 | 0.90592567 | 1.02843534 |
| KIR3DL2 | 0.778205 | 0.90592567 | 1.02909338 |
| MRC1 | 0.777206 | 0.90592567 | 1.17974986 |
| APP | 0.787 | 0.90713791 | 1.03746308 |
| IRF4 | 0.7835 | 0.90713791 | 1.03826556 |
| LCP2 | 0.785833 | 0.90713791 | 1.09603206 |
| LILRB5 | 0.781927 | 0.90713791 | 0.99227114 |
| MX1 | 0.788169 | 0.90713791 | 1.09320427 |
| PRDM1 | 0.788169 | 0.90713791 | 1.04931054 |
| PLA2G2E | 0.800459 | 0.919548 | 1.00293577 |
| B3GAT1 | 0.807183 | 0.9203395 | 0.98248658 |
| CUL9 | 0.806889 | 0.9203395 | 1.19257914 |
| IKZF3 | 0.80457 | 0.9203395 | 0.91413174 |
| LOC389386 | 0.80457 | 0.9203395 | 0.97716494 |
| BATF2 | 0.809272 | 0.92099985 | 1.08594497 |
| STAT5A | 0.817517 | 0.9286506 | 1.05717939 |
| GBP6 | 0.821601 | 0.93155504 | 0.89799605 |
| TBK1 | 0.823419 | 0.93188421 | 1.12946847 |
| CD8A | 0.828149 | 0.93204961 | 0.968047 |
| CTLA4_all | 0.825783 | 0.93204961 | 1.11452424 |
| TNFSF4 | 0.826835 | 0.93204961 | 1.12303898 |
| PSMD7 | 0.832884 | 0.93393243 | 0.98874898 |
| TAPBP | 0.832884 | 0.93393243 | 0.98977836 |
| PDGFRB | 0.83849 | 0.9384593 | 0.97786716 |
| PTGER4 | 0.839998 | 0.9384593 | 1.06076472 |
| PSMB7 | 0.856647 | 0.95356692 | 1.00243227 |
| TNFRSF14 | 0.855455 | 0.95356692 | 1.02925253 |
| ITGAX | 0.868579 | 0.96508778 | 1.04980553 |
| CD45R0 | 0.87336 | 0.96512609 | 1.02663383 |
| FCGR2A.C | 0.870969 | 0.96512609 | 1.04707472 |
| ZEB1 | 0.872163 | 0.96512609 | 1.10315843 |
| IL18 | 0.890158 | 0.98191027 | 1.01696117 |
| GZMB | 0.897333 | 0.98803814 | 0.98751913 |
| ITGB1 | 0.899735 | 0.98889793 | 1.02570048 |
| TAL1 | 0.902935 | 0.99063013 | 0.96736839 |
| CCL24 | 0.906946 | 0.99146427 | 1.16864602 |
| TNFRSF17 | 0.90635 | 0.99146427 | 1.01929115 |

| | | | |
|---|---|---|---|
| EGR1 | 0.911509 | 0.99447646 | 0.97408191 |
| KLRD1 | 0.912962 | 0.99447646 | 1.13724572 |
| BCL10 | 0.981834 | 1 | 0.99146668 |
| C1R | 1 | 1 | 1 |
| C1S | 1 | 1 | 1 |
| C7 | 1 | 1 | 1 |
| C8A | 1 | 1 | 1 |
| C8B | 1 | 1 | 1 |
| C9 | 1 | 1 | 1 |
| CCL11 | 1 | 1 | 1 |
| CCL16 | 1 | 1 | 1 |
| CCL26 | 1 | 1 | 1 |
| CCR8 | 1 | 1 | 1 |
| CCRL1 | 0.993459 | 1 | 0.9769923 |
| CD1A | 1 | 1 | 1 |
| CD34 | 1 | 1 | 1 |
| CD48 | 0.946747 | 1 | 1.05431641 |
| CDH5 | 1 | 1 | 1 |
| CFI | 1 | 1 | 1 |
| CLU | 1 | 1 | 1 |
| CXCL12 | 0.976741 | 1 | 0.97463052 |
| DEFB103B | 1 | 1 | 1 |
| GBP1 | 0.950373 | 1 | 1.02038819 |
| HAMP | 1 | 1 | 1 |
| HFE | 1 | 1 | 1 |
| IFNA2 | 1 | 1 | 1 |
| IFNB1 | 1 | 1 | 1 |
| IFNGR1 | 0.929841 | 1 | 1.01551595 |
| IL17A | 1 | 1 | 1 |
| IL17B | 1 | 1 | 1 |
| IL18R1 | 0.956418 | 1 | 0.99622314 |
| IL21 | 1 | 1 | 1 |
| IL22RA2 | 1 | 1 | 1 |
| IL27 | 0.996179 | 1 | 1.05060234 |
| IL29 | 1 | 1 | 1 |
| IL2RB | 0.952791 | 1 | 1.01714186 |
| IL3 | 0.993459 | 1 | 0.98962627 |
| IL5 | 1 | 1 | 1 |
| IL9 | 1 | 1 | 1 |
| IRAK4 | 0.956418 | 1 | 1.07324764 |
| ITGA5 | 0.943122 | 1 | 1.01115958 |
| ITLN2 | 1 | 1 | 1 |
| KIR3DL3 | 1 | 1 | 1 |

| | | | |
|---|---|---|---|
| KLRAP1 | 0.951339 | 1 | 0.9944749 |
| MAPKAPK2 | 0.969727 | 1 | 1.06804727 |
| MASP1 | 1 | 1 | 1 |
| MBL2 | 1 | 1 | 1 |
| MUC1 | 1 | 1 | 1.03402667 |
| MYD88 | 0.996366 | 1 | 0.99793393 |
| NOS2 | 0.993459 | 1 | 0.97639668 |
| TLR7 | 0.979226 | 1 | 1.01148386 |
| TNFSF11 | 1 | 1 | 1.01017007 |

**Supplementary Table 7**. Comparison between Null and IL-1b-induced genes, showing p-value, q-value and fold change.

| | Pre-Tx (V1) | | | Post-Tx (V2) | | |
|---|---|---|---|---|---|---|
| **Gene** | **p-value** | **q-value** | **Fold change** | **p-value** | **q-value** | **Fold change** |
| RELB | 8.61E-12 | 9.21E-10 | 4.89 | 0.0768 | 0.29 | 1.34 |
| ICAM1 | 1.11E-10 | 1.98E-09 | 6.52 | 0.0719 | 0.29 | 1.56 |
| IRAK3 | 1.11E-10 | 1.98E-09 | 6.58 | 0.0282 | 0.29 | 1.68 |
| NFKB2 | 8.59E-11 | 1.98E-09 | 4.06 | 0.0548 | 0.29 | 1.52 |
| NFKBIA | 1.11E-10 | 1.98E-09 | 5.83 | 0.0383 | 0.29 | 1.57 |
| TNFAIP3 | 3.84E-11 | 1.98E-09 | 5.52 | 0.0241 | 0.29 | 1.58 |
| NFKB1 | 1.82E-10 | 2.79E-09 | 3.84 | 0.0819 | 0.29 | 1.53 |
| ATG7 | 2.32E-10 | 3.11E-09 | 4.74 | 0.0719 | 0.29 | 1.47 |
| CLEC5A | 4.66E-10 | 5.54E-09 | 8.31 | 0.0930 | 0.29 | 1.72 |
| CDKN1A | 5.83E-10 | 6.24E-09 | 8.82 | 0.0629 | 0.29 | 1.88 |
| BCL3 | 1.12E-09 | 9.21E-09 | 5.82 | 0.1957 | 0.45 | 1.26 |
| CCRL2 | 1.12E-09 | 9.21E-09 | 10.45 | 0.0305 | 0.29 | 2.41 |
| IL8 | 1.12E-09 | 9.21E-09 | 9.53 | 0.0261 | 0.29 | 2.24 |
| NFKBIZ | 1.38E-09 | 1.06E-08 | 4.79 | 0.0548 | 0.29 | 1.53 |
| SRC | 2.55E-09 | 1.82E-08 | 4.62 | 0.0873 | 0.29 | 1.61 |
| TICAM1 | 4.58E-09 | 3.06E-08 | 4.21 | 0.2060 | 0.45 | 1.42 |
| BATF | 5.54E-09 | 3.29E-08 | 2.50 | 0.6751 | 0.87 | 0.99 |
| IL1RN | 5.54E-09 | 3.29E-08 | 9.93 | 0.0873 | 0.29 | 2.40 |
| IRAK2 | 6.68E-09 | 3.57E-08 | 9.63 | 0.0412 | 0.29 | 2.30 |
| JAK3 | 6.68E-09 | 3.57E-08 | 2.42 | 1.0000 | 1.00 | 1.02 |
| CTSS | 9.65E-09 | 4.49E-08 | 3.33 | 0.5340 | 0.78 | 1.17 |
| PTGS2 | 9.65E-09 | 4.49E-08 | 18.16 | 0.0768 | 0.29 | 3.38 |
| TNFAIP6 | 9.65E-09 | 4.49E-08 | 5.42 | 0.0511 | 0.29 | 1.83 |
| CXCL2 | 1.38E-08 | 6.17E-08 | 12.59 | 0.0412 | 0.29 | 2.29 |
| MARCO | 2.54E-08 | 1.09E-07 | 20.91 | 0.4004 | 0.70 | 1.46 |
| BID | 2.77E-08 | 1.14E-07 | 2.99 | 0.2591 | 0.52 | 1.75 |
| CCL20 | 3.27E-08 | 1.30E-07 | 13.80 | 0.0930 | 0.29 | 2.11 |
| PLAU | 4.56E-08 | 1.68E-07 | 24.71 | 0.0099 | 0.29 | 6.23 |
| SOCS3 | 4.56E-08 | 1.68E-07 | 3.98 | 0.2765 | 0.55 | 1.36 |
| TRAF1 | 5.37E-08 | 1.91E-07 | 3.27 | 0.0873 | 0.29 | 1.41 |
| CXCL1 | 7.40E-08 | 2.55E-07 | 5.37 | 0.0930 | 0.29 | 1.70 |
| CD14 | 1.86E-07 | 6.23E-07 | 3.02 | 0.2514 | 0.52 | 1.17 |
| TNF | 2.89E-07 | 9.39E-07 | 3.22 | 0.0673 | 0.29 | 1.57 |
| CD46 | 5.11E-07 | 1.61E-06 | 2.79 | 0.9424 | 0.98 | 1.04 |
| CASP8 | 5.88E-07 | 1.80E-06 | 2.23 | 0.9654 | 0.98 | 1.01 |
| CD83 | 6.74E-07 | 1.95E-06 | 4.23 | 0.0819 | 0.29 | 1.70 |
| POU2F2 | 6.74E-07 | 1.95E-06 | 2.46 | 0.3314 | 0.62 | 1.26 |
| KCNJ2 | 8.85E-07 | 2.43E-06 | 3.93 | 0.1333 | 0.35 | 1.74 |
| TNFSF15 | 8.85E-07 | 2.43E-06 | 12.98 | 0.0511 | 0.29 | 2.63 |
| CSF3R | 2.50E-06 | 6.69E-06 | 3.60 | 0.6751 | 0.87 | 0.92 |
| IL1B | 3.63E-06 | 9.46E-06 | 5.83 | 0.1333 | 0.35 | 1.81 |

| | | | | | | |
|---|---|---|---|---|---|---|
| CCL3 | 4.62E-06 | 1.18E-05 | 6.01 | 0.0511 | 0.29 | 2.26 |
| FCER1A | 4.93E-06 | 1.23E-05 | 0.09 | 0.5102 | 0.78 | 1.80 |
| SLAMF7 | 5.21E-06 | 1.27E-05 | 3.80 | 0.1258 | 0.35 | 1.81 |
| LILRB4 | 5.87E-06 | 1.39E-05 | 2.64 | 0.3462 | 0.63 | 1.30 |
| CD274 | 6.60E-06 | 1.50E-05 | 5.95 | 0.0816 | 0.29 | 4.00 |
| FER1L3 | 6.60E-06 | 1.50E-05 | 4.14 | 0.8060 | 0.95 | 0.75 |
| S1PR1 | 9.34E-06 | 2.08E-05 | 0.34 | 0.7837 | 0.93 | 1.19 |
| CD36 | 1.17E-05 | 2.51E-05 | 2.98 | 0.2514 | 0.52 | 1.34 |
| CXCR2 | 1.17E-05 | 2.51E-05 | 3.61 | 0.6751 | 0.87 | 0.92 |
| CD244 | 2.81E-05 | 5.89E-05 | 1.93 | 0.9424 | 0.98 | 1.10 |
| ICAM5 | 3.93E-05 | 8.09E-05 | 12.63 | 0.0149 | 0.29 | 3.74 |
| C3 | 4.71E-05 | 9.52E-05 | 4.26 | 0.4648 | 0.75 | 1.35 |
| IL16 | 7.03E-05 | 0.0001 | 1.93 | 0.8738 | 0.97 | 0.98 |
| CCL7 | 0.0001 | 0.0002 | 2.68 | 0.6268 | 0.86 | 1.47 |
| CD40 | 0.0002 | 0.0003 | 1.75 | 0.1186 | 0.33 | 1.26 |
| CCND3 | 0.0002 | 0.0004 | 2.16 | 0.8151 | 0.95 | 0.94 |
| NFATC3 | 0.0002 | 0.0004 | 1.67 | 0.7837 | 0.93 | 1.02 |
| CYBB | 0.0002 | 0.0004 | 1.58 | 0.9195 | 0.98 | 1.02 |
| IKBKE | 0.0002 | 0.0004 | 1.56 | 0.5532 | 0.80 | 1.13 |
| IL6 | 0.0003 | 0.0006 | 4.94 | 0.0378 | 0.29 | 3.96 |
| CCL4 | 0.0004 | 0.0008 | 3.22 | 0.0383 | 0.29 | 2.15 |
| IFIT2 | 0.0007 | 0.0011 | 3.28 | 0.9650 | 0.98 | 1.32 |
| CASP2 | 0.0010 | 0.0017 | 1.59 | 0.8966 | 0.97 | 1.01 |
| IL7R | 0.0013 | 0.0021 | 1.46 | 0.5340 | 0.78 | 1.13 |
| IL1A | 0.0016 | 0.0026 | 3.41 | 0.1186 | 0.33 | 2.50 |
| LILRB1 | 0.0023 | 0.0036 | 1.69 | 0.1118 | 0.33 | 1.70 |
| CCL19 | 0.0028 | 0.0044 | 0.24 | 0.2195 | 0.47 | 1.62 |
| LAIR1 | 0.0032 | 0.0050 | 1.51 | 0.8966 | 0.97 | 0.95 |
| SELE | 0.0035 | 0.0053 | 2.58 | 0.1408 | 0.35 | 1.49 |
| SELL | 0.0046 | 0.0069 | 1.66 | 0.7616 | 0.93 | 0.86 |
| KLF2 | 0.0060 | 0.0089 | 1.68 | 0.6334 | 0.86 | 0.92 |
| CXCL13 | 0.0106 | 0.0155 | 0.36 | 0.4326 | 0.74 | 1.67 |
| IL2RA | 0.0135 | 0.0193 | 1.55 | 0.0871 | 0.29 | 2.76 |
| ITGA4 | 0.0133 | 0.0193 | 1.38 | 0.5728 | 0.81 | 1.34 |
| CD80 | 0.0216 | 0.0304 | 0.42 | 0.1483 | 0.36 | 1.84 |
| IL23A | 0.0220 | 0.0306 | 2.50 | 0.6928 | 0.87 | 2.00 |
| TNFSF12 | 0.0262 | 0.0359 | 1.48 | 0.3462 | 0.63 | 0.91 |
| LILRA3 | 0.0276 | 0.0374 | 1.57 | 0.8285 | 0.95 | 1.18 |
| LILRA1 | 0.0280 | 0.0375 | 0.31 | 0.4646 | 0.75 | 0.62 |
| CCL23 | 0.0344 | 0.0455 | 0.42 | 0.3250 | 0.62 | 1.50 |
| CD40LG | 0.0396 | 0.0517 | 1.48 | 0.6964 | 0.87 | 1.26 |
| ADA | 0.0437 | 0.0557 | 1.30 | 0.9885 | 1.00 | 0.93 |
| IDO1 | 0.0437 | 0.0557 | 1.34 | 0.1118 | 0.33 | 1.73 |

| | | | | | | |
|---|---|---|---|---|---|---|
| AHR | 0.0482 | 0.0607 | 1.28 | 0.5151 | 0.78 | 1.17 |
| DUSP4 | 0.0505 | 0.0628 | 3.28 | 0.0299 | 0.29 | 3.63 |
| PDCD1LG 2 | 0.1326 | 0.1630 | 2.27 | 0.5709 | 0.81 | 1.59 |
| CCL22 | 0.1470 | 0.1729 | 0.94 | 0.6541 | 0.87 | 1.44 |
| CD1D | 0.1462 | 0.1729 | 0.41 | 0.6253 | 0.86 | 0.76 |
| IL10 | 0.1470 | 0.1729 | 1.76 | 0.5193 | 0.78 | 1.23 |
| SIGIRR | 0.1426 | 0.1729 | 0.88 | 0.4604 | 0.75 | 0.87 |
| HLA.DMB | 0.1615 | 0.1878 | 1.23 | 0.7837 | 0.93 | 1.15 |
| CCL2 | 0.1660 | 0.1889 | 0.65 | 0.4965 | 0.78 | 1.59 |
| PYCARD | 0.1643 | 0.1889 | 0.87 | 0.8491 | 0.97 | 0.86 |
| ALDH1A1 | 0.1770 | 0.1993 | 1.51 | 0.8951 | 0.97 | 0.95 |
| HLA.DMA | 0.2209 | 0.2462 | 1.35 | 0.8738 | 0.97 | 0.94 |
| CCR2 | 0.2419 | 0.2668 | 0.46 | 0.2060 | 0.45 | 0.81 |
| SELPLG | 0.3421 | 0.3735 | 0.60 | 0.3570 | 0.64 | 0.66 |
| TNFRSF9 | 0.4347 | 0.4698 | 1.29 | 0.1360 | 0.35 | 2.14 |
| LAMP3 | 0.4412 | 0.4721 | 0.44 | 0.0930 | 0.29 | 2.19 |
| BATF3 | 0.4652 | 0.4928 | 1.06 | 0.4384 | 0.74 | 1.59 |
| CISH | 0.4988 | 0.5233 | 1.48 | 0.1543 | 0.36 | 2.55 |
| EBI3 | 0.6138 | 0.6376 | 1.38 | 0.3287 | 0.62 | 1.83 |
| MSR1 | 0.6240 | 0.6420 | 0.87 | 0.4733 | 0.76 | 1.50 |
| GAS6 | 0.6462 | 0.6585 | 1.02 | 0.9509 | 0.98 | 0.97 |
| TNFRSF4 | 0.9430 | 0.9519 | 1.03 | 0.1558 | 0.36 | 1.93 |
| CYB561 | 0.9601 | 0.9601 | 0.82 | 0.6823 | 0.87 | 1.10 |

**Supplementary Table 8**. List of differentially expressed genes between LTBI and TB that are induced by IL-1b, with their respective p and q values, as well as fold-change.
